# Multi-scale model of axonal and dendritic polarization by transcranial direct current stimulation in realistic head geometry

**DOI:** 10.1101/2023.08.23.554447

**Authors:** Aman S. Aberra, Ruochen Wang, Warren M. Grill, Angel V. Peterchev

## Abstract

1.

**Background:** Transcranial direct current stimulation (tDCS) is a non-invasive brain stimulation modality that can alter cortical excitability. However, it remains unclear how the subcellular elements of different neuron types are polarized by specific electric field (E-field) distributions.

**Objective:** To quantify neuronal polarization generated by tDCS in a multi-scale computational model.

**Methods:** We embedded layer-specific, morphologically-realistic cortical neuron models in a finite element model of the E-field in a human head and simulated steady-state polarization generated by conventional primary-motor-cortex–supraorbital (M1–SO) and 4×1 high-definition (HD) tDCS. We quantified somatic, axonal, and dendritic polarization of excitatory pyramidal cells in layers 2/3, 5, and 6, as well as inhibitory interneurons in layers 1 and 4 of the hand knob.

**Results:** Axonal and dendritic terminals were polarized more than the soma in all neurons, with peak axonal and dendritic polarization of 0.92 mV and 0.21 mV, respectively, compared to peak somatic polarization of 0.07 mV for 1.8 mA M1–SO stimulation. Both montages generated regions of depolarization and hyperpolarization beneath the M1 anode; M1–SO produced slightly stronger, more diffuse polarization peaking in the central sulcus, while 4×1 HD produced higher peak polarization in the gyral crown. Simulating polarization by uniform local E-field approximated the spatial distribution of tDCS polarization but produced large errors in some regions.

**Conclusions:** Polarization of pre- and postsynaptic compartments of excitatory and inhibitory cortical neurons may play a significant role in tDCS neuromodulation. These effects cannot be predicted from the E-field distribution alone but rather require calculation of the neuronal response.

## 2. Introduction

Transcranial direct current stimulation (tDCS) is a non-invasive technique to modulate brain activity by applying weak electrical currents (typically 1–2 mA) via electrodes placed on the scalp. There is widespread interest in tDCS as an inexpensive, well-tolerated tool for neuroscience research and as a potential therapy for numerous psychiatric and neurological disorders [1–3]. Unfortunately, the outcomes of tDCS routinely suffer from high variability and low efficacy [4–7]. Clarifying the neural mechanisms of tDCS is critical to optimizing it as a tool for targeted neuromodulation. Using biophysically-realistic neuron models combined with anatomically-detailed head models, we simulated the direct polarization generated by common tDCS electrode montages within different cortical layers, morphological variants, and subcellular domains.

tDCS applied to the hand region of motor cortex (M1) can facilitate or suppress cortical excitability, and effects can last up to 60 minutes after stimulation depending on stimulus duration and polarity [8,9]. These neuromodulatory after-effects are thought to be mediated by subthreshold polarization of cortical neurons [10]. Subthreshold shifts in membrane potential alter endogenous activity *in vitro*, e.g., through shifts in spike timing or rate [11,12] or in the amplitude of post-synaptic potentials [13,14], and affect induction of synaptic plasticity [15,16]. However, the mechanisms of both the acute and long-lasting effects of polarization are still unclear, particularly when considering the complex cortical geometry in humans and morphological and electrophysiological diversity of cortical neurons.

Most modeling and *in vitro* experiments investigating the effects of dc electric fields on neurons applied spatially uniform E-fields, motivated by the quasi-uniform assumption [17]. This assumption treats the local E-field as uniform, since the large, distant current sources at the scalp generate an E-field in the brain with low spatial gradient at the scale of individual neurons. Using this approximation, stimulation with a uniform dc E-field establishes a gradient of polarization across a neuron’s morphology, with depolarization at the anodal end, hyperpolarization at the cathodal end, and a zero-point near the center [18,19]. Peak polarization occurs at terminations, leading to strongest polarization at the most distal basal and apical dendrites and at axon terminations [20,21]. Somatic polarization depends on the electrotonic distance to either end and the complex interplay of current flow in the numerous dendritic and axonal branches where direct E-field induced polarization occurs. In line with these theoretical principles, pyramidal cells’ longer somatodendritic extent leads to higher somatic polarization *in vitro* (∼0.25 mV/V/m), while more radially symmetric interneurons experience, on average, negligible somatic polarization [22,23]. The electrotonic position of the pyramidal cell soma leads to somatic depolarization when the anode is at the pial end and somatic hyperpolarization when the cathode is at its pial end. These findings led to a focus on the expected direct effects on pyramidal cell somas when designing tDCS experiments and clinical protocols, as tDCS is expected to be soma-depolarizing underneath the anode and soma-hyperpolarizing underneath the cathode.

However, polarization in the axonal and dendritic terminals is expected to be several times higher than in the soma. Axon terminals are polarized at least 2 – 6× more than the soma in uniform E-field [13,20,21,24], and axon terminal polarization may affect the magnitude or timing of pre-synaptic release after endogenous spike initiation [21]. Dendritic polarization can also alter input–output properties in a pathway specific manner [13,14,25]. Polarization in highly branched axonal and dendritic arbors has a complex dependence on E-field orientation, making challenging the extrapolation of findings in brain slices with uniform E-field to tDCS in humans. Finally, it has not been established how well the quasi-uniform assumption holds when considering the non-uniform E-field distribution generated by tDCS along spatially extended morphologies of cortical neurons. Therefore, characterizing the effects of tDCS-generated E-fields on pre- and post-synaptic compartments of both excitatory and inhibitory cell types is important for a comprehensive understanding of how tDCS modulates cortical circuits.

Simulating the tDCS-generated E-field distribution in anatomically-accurate head models is increasingly common and intended to ensure current is delivered to the desired target while accounting for individual anatomy [26,27]. Some investigators suggested E-field components normal and tangential to the cortical surface can be used as proxies for the direct neural effects. For example, the normal component was used in multi-electrode optimization [28] and in analyzing neuromodulatory after-effects of tDCS [29–31]. However, it remains unclear how well these macroscopic E-field components can determine polarization of different neural elements, accounting for their diverse orientations and morphologies.

Here, we report a multi-scale model of the direct neural effects of tDCS that combines a finite element method (FEM) simulation of the E-field in an magnetic resonance imaging (MRI) MRI-derived head model with biophysically realistic multi-compartmental models of cortical neurons [20,32]. Critically, the neuron models include both realistic dendritic and axonal morphologies with voltage-gated ion channels, excitatory and inhibitory cell types in multiple cortical layers, and multiple morphological variants of each cell type. We quantified the polarization of all subcellular compartments (soma, axons, and dendrites) by both uniform E-fields and E-field distributions generated by conventional 7×5 cm rectangular-pad tDCS and the more focal 4×1 high-definition (HD) tDCS in a patch of cortex surrounding the hand knob. Recently, Chung et al. coupled a subset of these neuron models to an E-field head model to simulate activation thresholds and polarization distributions for similar tES electrode montages [33]; however, their quantitative analysis focused on the effect of morphology on suprathreshold responses. In contrast, we evaluated how well the quasi-uniform approximation and decomposition of the macroscopic E-field into normal and tangential components can be used to estimate the polarization responses of the detailed model neurons.

## 3. Methods

We used our previously developed framework [32] to embed populations of biophysically-realistic neuron models in cortical regions of MRI-derived head models and simulate their response in NEURON [34]. We quantified the steady-state polarization generated in all neuronal compartments of layer-specific excitatory and inhibitory cell types by two tDCS electrode montages targeting the motor hand knob region and by stimulation with uniform electric field applied at the full range of possible directions.

### 3.1. Volume conductor model of electric field

#### 3.1.1. Head model

Computation of electric fields (E-fields) generated by scalp electrodes was performed in SimNIBS v3.0 using the finite element method (FEM) [35]. The head model was generated from T1 and T2-weighted MR images example data set (*almi5*) included with SimNIBS using the *mri2mesh* pipeline [36], which output a mesh with 1.6 million vertices and 8.8 million tetrahedra segmented into five tissues: white matter, gray matter, cerebrospinal fluid, bone, and scalp tissue. The white matter layer was assigned default anisotropic conductivity settings using the included diffusion tensor imaging (DTI) data and the volume normalized approach, with the mean conductivity of each tensor scaled to match isotropic conductivity of 0.126 S/m [30]. The maximum eigenvalue and ratio between eigenvalues for the conductivity tensors was 10 and 2, respectively. The remaining four tissue volumes were assigned isotropic conductivities [30] (in S/m): gray matter: 0.275, cerebrospinal fluid: 1.654, bone: 0.010, scalp: 0.465.

#### 3.1.2. Electrodes

We simulated two electrode montages [37] using the built-in tools within SimNIBS for electrode generation and meshing: 1) conventional 7×5 cm^2^ rectangular pad M1–SO configuration (Figure 1A, top) and 2) 4×1 HD ring configuration, with four disk electrodes surrounding a central electrode centered over the M1 hand knob (Figure 1A, bottom). For the M1–SO configuration, we centered one pad above the M1 hand knob [38] and the other above the contralateral eye (supraorbital) region. The pad electrodes modeled a silicone rubber electrode inside a sponge soaked in saline, with a 1 mm silicone rubber electrode layer between 1 mm thick “sponge” layers, assigned conductivities of 29 and 1.4 S/m, respectively [37,39]. For anodal M1–SO stimulation, we applied 1 mA current to the M1 electrode and −1 mA to the SO electrode. For the 4×1 HD configuration, we used disk electrodes with 8 mm diameter arranged with the four surround electrodes placed with a center-to-center distance of 3 cm from the central electrode at 90° intervals. The disk electrodes consisted of a 1 mm thick rubber electrode above a 2 mm thick layer of saline gel, using the same conductivities as above. For anodal 4×1 HD stimulation, we computed the E-field distribution for 1 mA current applied to the central electrode and −0.25 mA to the four surround electrodes. Since all tissues were approximated as ohmic conductors, the E-field distributions at different stimulation intensities (with the same ratio of amplitudes in the 4×1 HD case) were obtained by linear scaling.

**Figure 1.**
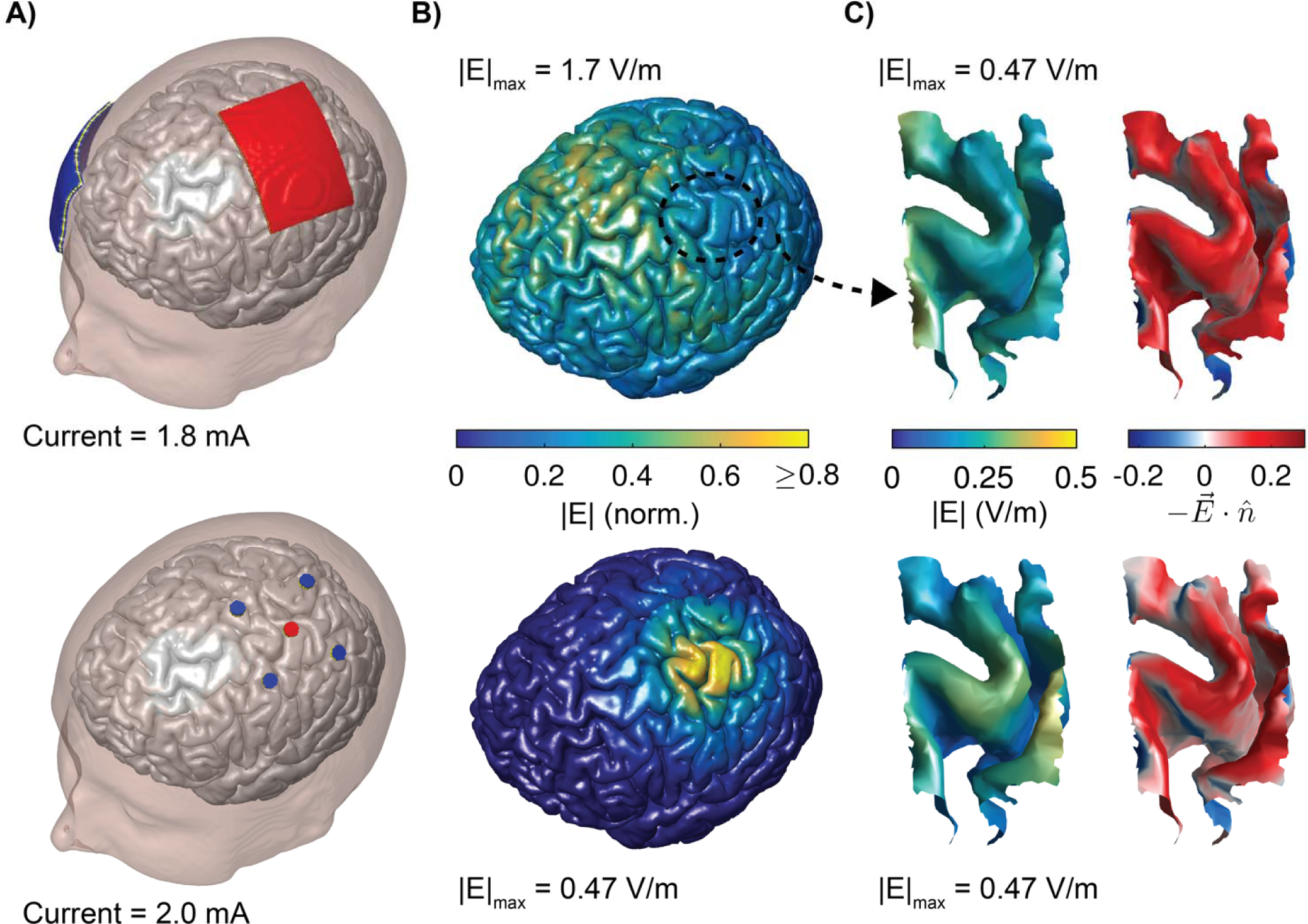
Finite element method (FEM) model of conventional M1–SO and 4×1 HD tDCS. A) Scalp and gray matter (GM) surfaces with anode in red, cathode in blue for conventional M1–SO montage with 7×5 cm rectangular pad electrodes (top) and HD 4×1 ring configuration centered over M1 hand knob (bottom). B) E-field magnitude distribution on full GM surface for M1–SO (top) and 4×1 HD (bottom) montages. C) E-field magnitude distribution on L5 surface mesh within region of interest (ROI) in M1 hand knob populated with neurons, where peak |E| was matched for the M1–SO (top) and 4×1 HD (bottom) montages (left). E-field component normal to cortical surface, indicating direction of current flow in the same region (right). Inward current is positive.

### 3.2. Neuron models

The details of the neuron models and method for placing them were described previously [32]. Briefly, we adapted a set of multi-compartmental, conductance-based cortical neuron models from the library of models released by the Blue Brain Project [40,41] in NEURON v7.7 [34]. These models included 3D, reconstructed dendritic and axonal morphologies from all 6 cortical layers with up to 13 different published, Hodgkin-Huxley-like ion channel models distributed in the soma, dendrites, and axon. The cell types included were: layer 1 neurogliaform cell (NGC) with dense axonal arborization, L2/3 pyramidal cell (PC), L4 large basket cell (LBC), L5 thick-tufted PC, and L6 tufted PC. The L1 NGC model exhibited burst non-accommodating firing, the L4 LBC exhibited continuous accommodating firing, and the PCs were all continuous adapting. The models were originally developed based on juvenile (P14) rat brain slice data, and to model mature, human neurons, we modified the models to account for age and species differences [20]. Each cell type had five virtual “clones” with stochastically varied dendritic and axonal geometries. The response of the models to subthreshold stimulation was previously validated based on *in vitro* recordings of somatic and axon polarization in response to uniform dc E-fields [20–22].

### 3.3. Embedding neurons in head model

To embed the layer-specific model neurons in the head model, we first extracted the precentral gyrus, central sulcus, and postcentral gyrus labeled regions generated by Freesurfer’s automatic cortical parcellation with the Destrieux Atlas [42]. We then cropped this region with a 31×24×50 mm box to define the region of interest (ROI) in which we embedded the model neuron populations. As described previously [32], we generated surface meshes representing each cortical layer to place and orient the model neurons. Here, we used layer depth boundaries from the recently published layer segmentation of the BigBrain histological atlas; in Von Economo area FA, the boundaries between adjacent layers were at normalized depths of 0.0993 (L1–L2/3), 0.466 (L2/3–L4), 0.524 (L4–L5), 0.753 (L5–L6) (total depth of gray matter is 1) [43]. Between these boundaries, we generated the cell placement surface meshes at the following normalized depths: L1: 0.01, L2/3: 0.4, L4: 0.5, L5: 0.75, L6: 0.95. The layer surface meshes were discretized with 3000 triangular elements per surface using the PyMeshLab 0.2 Python interface with MeshLab 2020.12 (Python v3.8) [44], resulting in surfaces with mean density of 2.4 elements per mm^2^. We also used MeshLab to improve the quality of the cell placement surface meshes by removing any self-intersecting faces, holes, and non-manifold edges and re-computing face normals. Single model neurons were placed in each of the 3000 elements in their respective layers, with their cell bodies centered in the triangle and somatodendritic axes aligned to the element normal. The model neurons were placed with random rotations about their somatodendritic axes in the azimuthal direction (see Figure 2A for definition of spherical coordinate system). However, this procedure still allows some axonal or dendritic branches to intersect with the pial surface, which is unrealistic, particularly in regions with thinner gray matter. We improved the robustness of our previous approach by detecting misplaced neuron morphologies using a fast algorithm for computing ray-triangle intersection [45]. Then, we determined the appropriate translation and azimuthal rotation to position the model neurons within anatomical boundaries using a binary search algorithm, which minimized deviation from their original position and retained alignment to the cortical columns. The maximal allowed translation was set based on the distance to the lower layer boundary from the soma, ensuring the cell bodies remained in their original layer.

**Figure 2.**
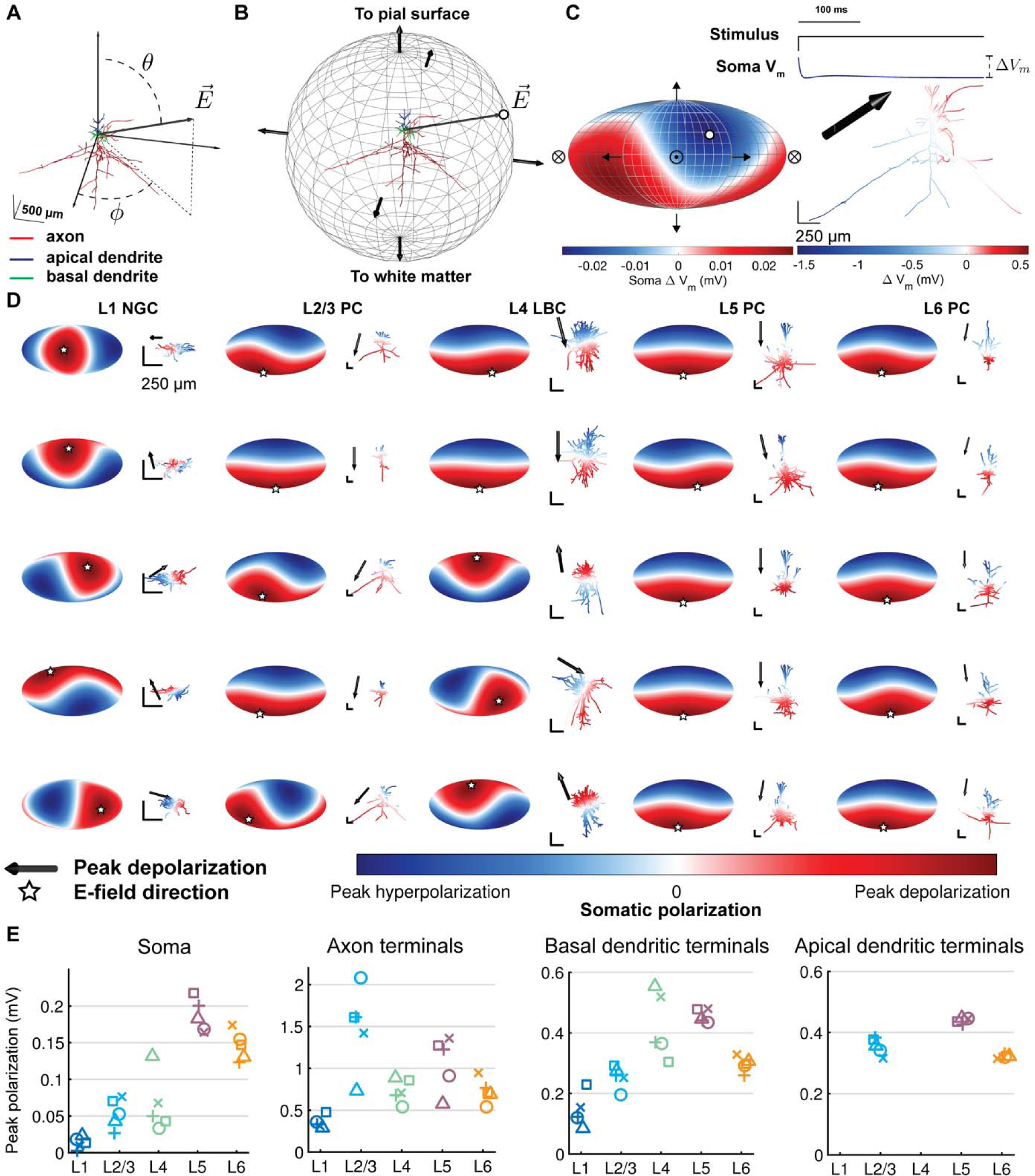
Dependence of polarization on cell type and direction of uniform E-field. A) Coordinate system shown for example model neuron (L2/3 PC). Somato-dendritic axis aligned to z-axis, with polar angle θ and azimuthal angle ϕ. B) Uniform E-field directions represented as normal vectors on sphere centered at origin. Steady-state polarization in a specific neural element (e.g., soma) was calculated for 266 directions spanning the sphere, and each polarization value was represented as a point on the sphere (white dot) corresponding to the E-field vector 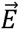. C) (left) 3D polarization–direction map projected into 2D using Mollweide projection. White dot indicates polarization value for example vector 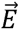 in B, crossed circle represents E-field pointing into the page, and circle with dot represents E-field pointing out of the page. (right, bottom) Color plot of membrane polarization at the end of a 500 ms 1 V/m uniform E-field (same direction as B). (right, top) Recording from soma demonstrating polarization ***ΔV_m_*** calculation. D) Polarization–direction maps of peak somatic polarization for all cell types and their virtual clones, i.e., models of the same cell type with stochastically varied morphologies. Clones of each cell type (columns) are ordered by peak somatic depolarization, with lowest peak depolarization at the bottom. Corresponding cell morphology plotted to the right of each map colored with steady-state polarization distribution for E-field direction producing peak somatic depolarization, indicated by black arrow and matching white star. All scale bars are 250 µm. E) Peak polarization for the 5 clones of each cell type, grouped by layer, in the soma, axon terminals, basal dendritic terminals, and apical dendritic terminals.

All five clones of each cell type were co-located within each element. Except where indicated otherwise above, mesh generation, placement of neuronal morphologies, extraction of voltages from the SimNIBS output, NEURON simulation control, analysis, and visualization were conducted in MATLAB (R2019a, The Mathworks, Inc., Natick, MA, USA).

### 3.4. Coupling electric fields to neuron models

#### 3.4.3. Spatial component

The quasi-static approximation allows separation of the tDCS-generated E-field into its spatial and temporal components [46,47]. This enables extraction of the spatial distribution of voltages from the FEM head model at a single current intensity and scaling over time by the stimulus waveform in the neuronal simulations. After placement of the model neuron populations and simulation of the E-field distribution generated by the tDCS surface electrodes, the voltages at the coordinates of each model neuron’s compartments were extracted using the SimNIBS functions for interpolation of field values within the FEM mesh (`get_fields_at_coordinates.m`). This function uses the 1^st^ order FEM basis functions of each tetrahedral element and voltages computed at the vertices to interpolate voltages at any coordinate within the corresponding element. The voltages were applied as extracellular voltages to the compartments of the cable neuron models using NEURON’s `extracellular` mechanism, comprising the spatial component of the extracellular E-field.

We also quantified the response of the model neurons to uniform E-fields by computing the extracellular potential *v_e_* at each compartment with position (x, y, z):

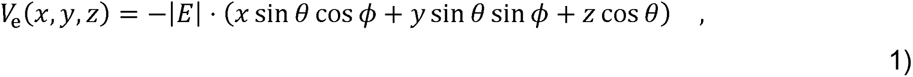

where the direction of the uniform E-field was given by polar angle θ and azimuthal angle ϕ, in spherical coordinates with respect to the somatodendritic axes (Figure 2A), and the potential at the origin (soma) was set to zero [20]. We applied 1 V/m uniform E-fields at directions spanning the polar and azimuthal directions in steps of 15°, totaling 266 directions. With this coordinate system, we defined downward E-field (120° < θ ≤ 180°), which is analogous to E-field directed inward relative to the cortical surface, upward E-field (0 ≤ θ ≤ 60°), analogous to E-field directed outward relative to the cortical surface, and transverse E-field (60° < θ ≤ 120), analogous to E-field tangential to the cortical surface.

#### 3.4.4. Temporal component

For the temporal E-field component in both cases, we uniformly scaled the distribution of potentials over time by a 500 ms rectangular pulse waveform. We quantified the steady-state polarization generated by tDCS by calculating the change in membrane voltage of each compartment at the end of the stimulus waveform relative to the initial state (shown in Figure 2C). As in our previous studies [20,32], the neuron models were discretized with isopotential compartments no longer than 20 µm and solved numerically using the backward Euler technique. However, we used a longer time step of 25 µs, rather than 5 µs, to facilitate longer simulation durations. The membrane potential was allowed to equilibrate to steady state before stimulation was applied. We focused our analysis on the distribution of compartmental polarizations within the soma, axons, apical dendrites, and basal dendrites. Similar to our approach with uniform E-field activation thresholds with TMS pulses [32], we generated 2D polarization–direction maps using the Mollweide projection (Figure 2B–C). While the somatic polarization can be described by a single value, we quantified polarization in the dendritic and axonal arbors using peak polarization, which is the maximum absolute value of polarization across the corresponding compartments. We retain the sign of polarization, allowing for this value to be either a depolarized or hyperpolarized compartment.

### 3.5. Method for estimating tDCS-generated polarization with uniform E-field simulations

In addition to using uniform E-field to characterize the sensitivity of each cell type and neural element to dc stimulation, we also implemented a method for estimating the polarization generated at any location within the non-uniform tDCS E-field distribution using the quasi-uniform assumption [17]. This assumption enables a simplified and rapid method of estimating compartmental polarizations in each neuron model relative to the full FEM-coupled simulations described above. In this method, for each model neuron in the head model, we extracted the E-field vector from the FEM solution at the soma and interpolated the polarization for a 1 V/m uniform E-field of the same direction from the polarization–direction map, after rotating the vector into the cell-centered coordinate system used in the uniform E-field simulations (Figure 2A). We then scaled this polarization value linearly by the amplitude of the E-field vector. This latter step relies on the separate assumption of linear membrane polarization, despite the presence of non-linear ion channels that may be activated at subthreshold membrane voltages. Therefore, the steady-state polarization Δ*V_m_* for a somatic E-field vector 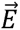 with polar and azimuthal components *E_θ_* and *E_ϕ_*, respectively, is given by:

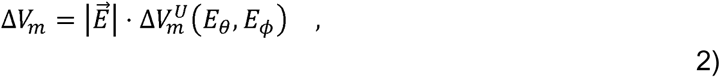

where 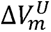 is the uniform E-field polarization interpolant derived from the polarization–direction map. The same procedure was carried out for peak basal dendritic, apical dendritic, and axonal polarization using the peak polarization–direction maps. The interpolant was implemented in MATLAB as a `griddedInterpolant` using linear interpolation. We quantified estimation error using this method with median absolute percent error (MedAPE), mean absolute percent error (MAPE), and mean absolute normalized error (MANE). MANE was calculated by normalizing absolute errors by the 0.975 quantile across the layer for each model neuron.

### 3.6. Code and data availability

The code and relevant data of this study will be made available on GitHub upon publication (https://github.com/Aman-A/tDCSsim_Aberra2023).

## 4. Results

### 4.1. Polarization sensitivity to electric field direction varies by cell type and element

Using 500 ms, 1 V/m uniform E-field pulses, we quantified the somatic polarization, as well as peak polarization in the apical dendritic, basal dendritic, and axonal compartments for excitatory and inhibitory cell types spanning the cortical layers across the full range of possible E-field directions. As shown previously [13,20,24,48], uniform E-fields applied with a dc pulse established a biphasic distribution of steady-state membrane polarization across the neuronal morphology, with depolarization at the cathodal end and hyperpolarization at the anodal end (Figure 2). Pyramidal cell somas were maximally depolarized with inward and hyperpolarized with outward E-fields consistently across clones (Figure 2D). In these cells, somatic polarization followed a smooth cosine dependency on the polar angle, with minimal variation due to azimuthal rotations of the E-field. In contrast, the sensitivity to E-field direction of polarization of the interneuron somas varied between clones. The L1 NGCs experienced maximal somatic depolarization for transverse (2/5 clones) or outward (3/5 clones) E-fields, and two of the L4 LBCs exhibited directional sensitivity similar to the pyramidal cells, while the others had either reversed polarization polarity or were depolarized and hyperpolarized most strongly with transverse E-fields. Peak somatic polarization across directions was highest in the L5 PCs (range: 0.16 – 0.22 mV) and L6 PCs (0.12 – 0.17 mV), followed by the L4 LBCs (0.033 – 0.13 mV), L2/3 PCs (0.027 – 0.076 mV), and L1 NGCs (0.003 – 0.023 mV). On average, peak somatic polarization was higher for inward and outward E-field directions than for transverse E-fields for all cell types, but only marginally for the L1 NGCs (Supplementary Figure S1).

Peak axonal and dendritic polarization was substantially higher than peak somatic polarization in all cells (Figure 2E) and across E-field directions, with the strongest polarization at the terminations. Therefore, peak axonal, basal dendritic, and apical dendritic *terminal* polarization was always synonymous with peak axonal, basal dendritic, and apical dendritic polarization, respectively, and subsequent analysis focused on terminal polarization.

Peak axon terminal polarization was highest in the L2/3 PCs (0.73 – 2.1 mV), followed by the L5 PCs (0.58 – 1.4 mV). The L6 PCs (0.54 – 0.95 mV) and L4 LBCs experienced similar peak axonal terminal polarization (0.54 – 0.89 mV), and the L1 NGCs experienced the lowest peak axonal terminal polarization (0.29 – 0.48 mV).

While the excitatory neurons had the highest axonal and somatic polarization, the inhibitory L4 LBCs experienced highest peak dendritic polarization (0.30 – 0.55 mV), followed by the L5 PCs (0.44 – 0.48 mV), L2/3 PCs (0.29 – 0.35 mV), L6 PCs (0.31 – 0.33 mV), and L1 NGCs (0.085 – 0.23 mV) (Figure 2E). The apical and basal dendrites of the pyramidal cells were polarized with opposite polarity and slightly weaker peak values (interneurons do not have an apical dendrite). Unlike somatic polarization, polarization–direction maps for peak axonal and dendritic polarization were more discontinuous and complex, exhibiting multiple local minima and maxima (Supplementary Figure S2– S4). These directions aligned with major axonal or dendritic branches specific to each clone’s arborization.

### 4.2. Polarization distributions

We simulated the response of the population of model neurons for 2.0 mA 4×1 HD tDCS, the upper limit of tolerable intensities for this montage [49], and 1.8 mA M1–SO tDCS with 7×5 cm rectangular pad electrodes, which we chose to generate identical peak E-field magnitude in the ROI of 0.47 V/m (Figure 1C). As expected, the 4×1 HD montage produced substantially more focal E-fields with peak E-field magnitude in the superficial precentral gyrus, while the M1–SO montage produced peak E-field (1.7 V/m) between the anode and cathode in a sulcal fold outside the ROI (Figure 1B). Figure 3 shows the spatial distribution of steady-state membrane polarization in the soma, axon, basal dendrites, and apical dendrites generated by both montages in each cortical layer, demonstrating regions of depolarization and hyperpolarization beneath the anode for both montages. The regions of depolarization and hyperpolarization for peak somatic, axonal, and basal dendritic compartments largely overlapped with the regions of inward and outward E-field component normal to the cortical surface, respectively (Figure 1C). The 4×1 HD montage generated stronger polarization in the gyral crown compared to the M1–SO montage, which is more clearly seen when visualizing the polarization differences between montages (Supplementary Figure S5). The distributions of polarization in the soma (Figure 3A) and basal dendrites (Figure 3C) were most similar, due to the small electrotonic distance between these compartments. Peak axonal polarization also had patterns of depolarization and hyperpolarization similar to those of the soma and basal dendrites, with peak depolarization in regions with inward E-field (positive normal component) and peak hyperpolarization in regions with outward E-field (negative normal component) (Figure 3B). Peak apical dendritic polarization in the PCs was always opposite in polarity to somatic polarization and peak basal dendritic polarization (Figure 3D).

**Figure 3.**
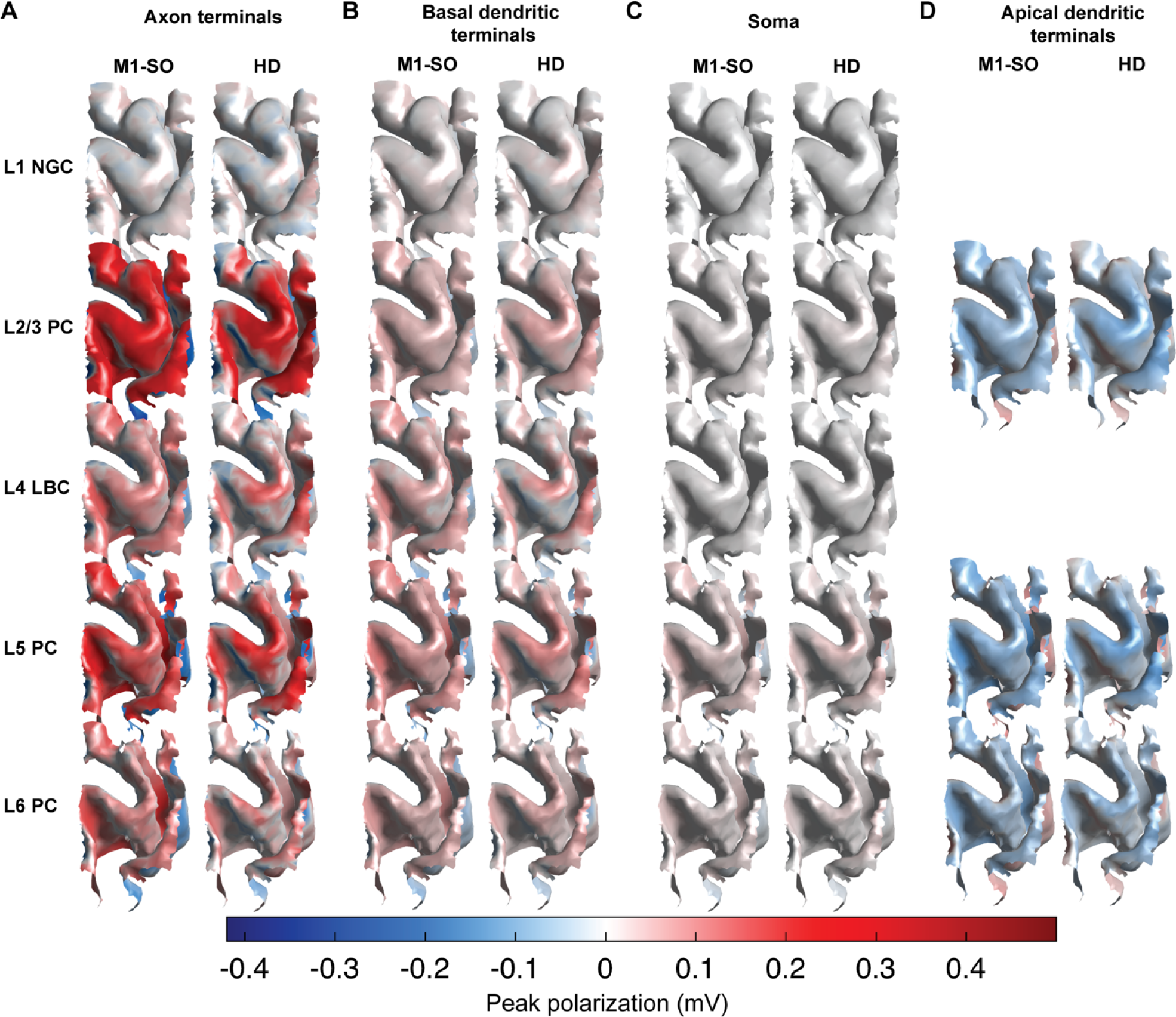
Spatial distribution of transmembrane polarization in hand knob ROI with M1–SO 7×5 cm rectangular pad +1.8 mA and 4×1 HD tDCS with +2.0 mA current through the target electrode. Surface plots are colored by median of peak polarization across clones at each position within A) axon terminals B) basal dendritic terminals, C) soma, and D) apical dendritic terminals for 1.8 mA M1–SO rectangular pad (left) and 2.0 mA 4×1 HD tDCS.

Figure 4 shows the distributions of compartmental polarizations in the simulated ROI for both montages. As with the uniform E-fields, all neurons experienced substantially higher peak axonal and dendritic polarizations than somatic polarization, with greatest somatic polarization in the L5 and L6 PCs, greatest peak axonal polarization in the L2/3 PCs, and greatest peak dendritic polarization in the dendrites of the L4 LBCs. Peak apical dendritic polarization in the PCs was slightly weaker than basal dendritic polarization, with strongest effects in the L5 PCs.

**Figure 4.**
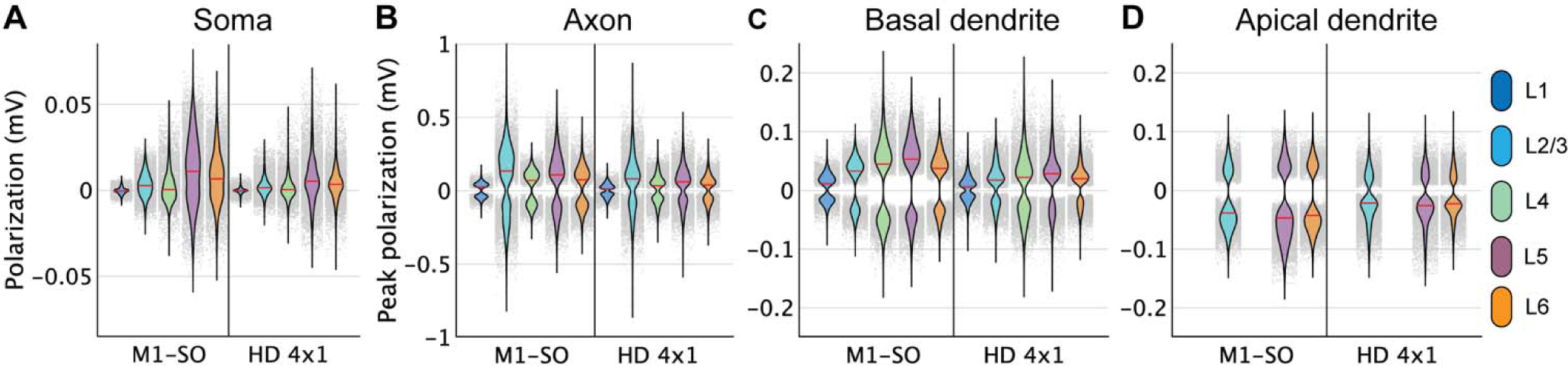
Distributions of polarization in ROI with M1–SO 7×5 cm rectangular pad at +1.8 mA and 4×1 HD tDCS at +2.0 mA. Each violin plot depicts the probability density estimate using a gaussian kernel of the A) somatic polarization, or peak polarization in B) axon terminals, C) basal dendritic terminals, and D) apical dendritic terminals of each layer for both electrode montages. Probability density was estimated using a normal kernel smoothing function with optimal bandwidth determined separately for each distribution. Point spread of polarization values plotted in gray, with each point representing the polarization value from a single model neuron. Note the different y-axis ranges.

Across the three atlas-defined cortical regions within the ROI, the 4×1 HD montage generated stronger peak, but lower median, polarization in the precentral gyrus than the M1–SO montage (Supplementary Figure S6). The M1–SO electrodes generated stronger polarization in the central sulcus and weaker polarization in the postcentral gyrus, based on both median and range, and all these responses follow from the E-field distributions. These trends were preserved across compartments, indicating differences in polarization largely followed differences in the macroscopic E-field distribution. The distributions of peak axonal and dendritic polarization were strongly bimodal, with strong depolarization and hyperpolarization, while the somatic polarization was more unimodal, both within the full ROI and within each of the three sub-regions (Figure 4, Supplementary Figure S6).

### 4.3. Correlation of polarization with E-field components varies by cell and compartment

We asked how well the somatic, axonal, or dendritic polarization were explained by the magnitude |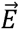|, normal component *E_n_*, or tangential component *E_t_* of the local E-field, extracted at each cell’s soma throughout the ROI. Focusing this analysis on the 7×5 cm M1–SO montage, |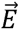| correlated weakly with the polarization in all compartments (*R*^2^ < 0.13) (not shown), and compartment-specific correlations improved when |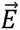| was regressed against absolute polarization values (*R*^2^: 0.12 – 0.61) (Figure 5). *E_n_* was more strongly correlated with somatic polarization than the |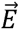| for all cells, but the correlation coefficients were substantially higher for the PCs (*R*^2^: 0.93 – 0.98) than the interneurons (*R*^2^: 0.39 – 0.41) (Figure 5A; Figure 6A). Compared to somatic polarization, peak axonal and dendritic terminal polarization had stronger correlations with |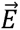|, but still with modest correlation coefficients (axonal *R*^2^: 0.36 – 0.51, dendritic *R*^2^: 0.48 – 0.61) (Figure 5B–D; Figure 6B–D). Peak axonal and dendritic terminal polarization also had higher correlations with *E_n_* than |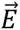| within cell type, with the exception of the L1 NGCs, which had negligible correlation of peak axonal terminal polarization with *E_n_* (*R*^2^ = 0.04) and low correlation with peak basal dendritic polarization (*R*^2^ = 0.34) (Figure 6). However, relative to somatic polarization, *E_n_* explained less variance in axonal and basal dendritic terminal polarization (axonal *R*^2^: 0.55 – 0.87, dendritic *R*^2^: 0.52 – 0.81; excluding L1 NGCs) (Figure 5; Figure 6). Apical dendritic polarization in the PCs had relatively high correlation with *E_n_* (*R*^2^: 0.79 – 0.92). Finally, *E_t_* had low correlations with polarization for all compartments and cell types (*R*^2^: 0.03 – 0.15) (Figure 5). Thus, the normal component of the E-field broadly captured the polarity and distribution of membrane polarization in the excitatory pyramidal cells, with the strongest correlations for somatic polarization. However, for somatic, axonal, and dendritic polarization of interneurons, as well as axonal polarization in the pyramidal cells, the normal component left substantial unexplained variance.

**Figure 5.**
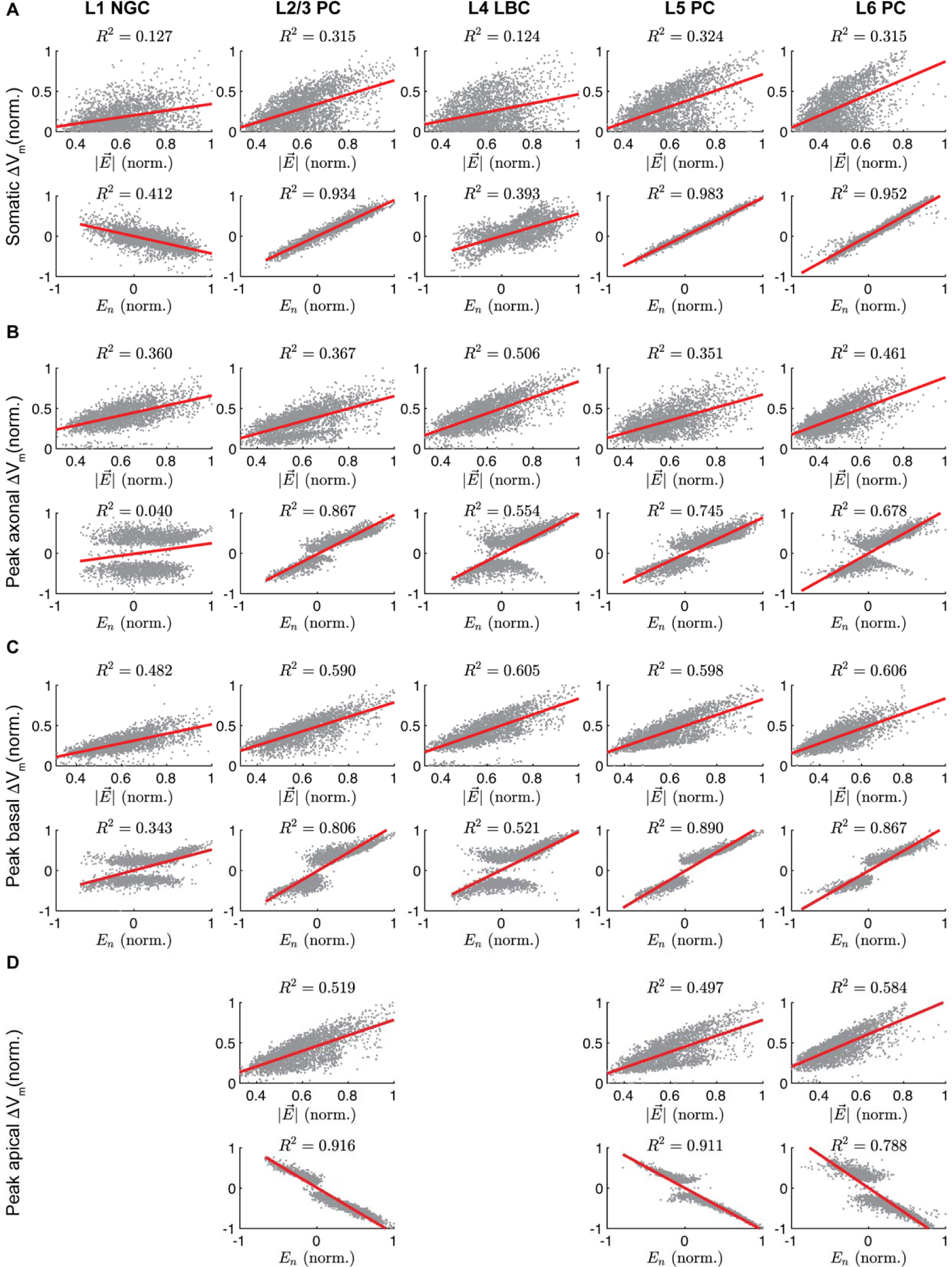
Correlation between membrane polarization and magnitude or component normal to pial surface of E-field vector at soma for M1–SO 7×5 cm rectangular pad montage. Normalized E-field components at each position throughout ROI plotted against compartmental polarizations with linear regressions (red) for median polarization across clones in A) soma, or median across clones of peak polarization in B) axon terminals, C) basal dendritic terminals, or D) apical dendritic terminals across clones. For E-field magnitude, absolute polarization was used.

**Figure 6.**
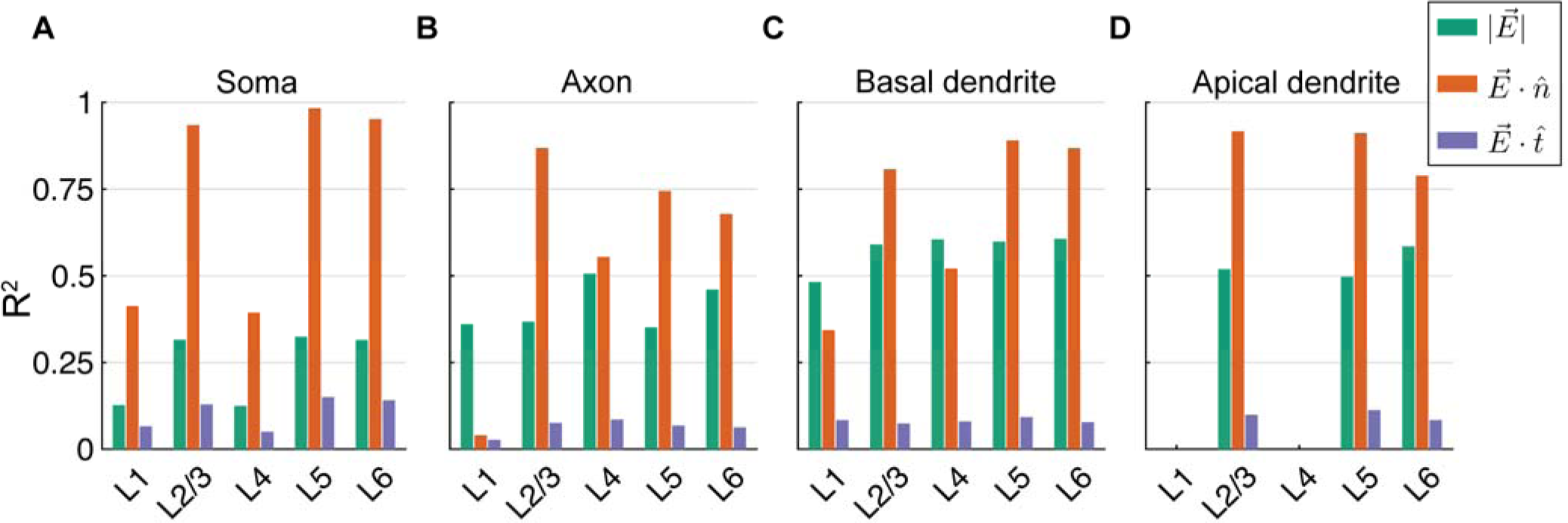
Coefficients of determination R^2^ between polarization and magnitude, normal, or tangential component of E-field vector at the soma for M1–SO 7×5 cm rectangular pad montage. Correlation for linear regressions of E-field components with A) median somatic polarization across clones, B) median of peak axon terminal polarization across clones, C) median of peak basal dendritic terminal polarization across clones, and D) median of peak apical dendritic terminal polarization across clones. Regression lines and underlying data shown in Figure 5. All correlation coefficients were statistically significant (p < 0.001).

### 4.4. Estimating polarization with uniform E-field simulations

The correlation analysis indicated that distributions of compartmental polarization were in some cases strongly correlated with components of the E-field extracted at the cell body across the ROI. We tested whether uniform E-field simulations can estimate rapidly the spatial distribution of steady-state membrane polarization generated by the non-uniform E-field computed in the FEM head model. In this method, we treated the E-field for a model neuron at a given location as uniform in magnitude and direction as an E-field vector from a specified reference point. Choosing the soma as this reference point, we extracted somatic E-field vector 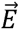 for each model neuron in the ROI, interpolated the somatic, peak axonal, peak basal dendritic, or peak apical dendritic polarization from the corresponding polarization–direction map (generated with 1 V/m uniform E-field stimulation), and scaled this polarization by |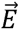|. This approach largely captured the spatial distribution of polarization, but with some regions of high relative error (Figure 7). The distributions of estimated polarization are depicted in Supplementary Figure S7 and error statistics across layers are listed in Supplementary Theable 1. Estimation error for all layers was slightly higher for the 4×1 M1 HD montage than the M1–SO montage, likely due to the higher E-field gradients. Mean absolute percent error (MAPE) was significantly higher in all compartments than the median, driven by outliers with high error mostly for polarization values close to zero. While estimation error using these relative error metrics was highest for somatic polarization, MANE for each model neuron was highest for peak axonal polarization. Between layers, estimation errors for somatic polarization were highest in the L1 NGCs (MedAPE: 20.31% M1–SO and 20.21% 4×1 HD) and L6 PCs (MedAPE: 28.89% M1–SO and 31.61% 4×1 M1-HD) and below 11% in L2–4 for both montages (Figure 8). This trend between layers held for axonal and dendritic polarization, with slightly higher errors for axonal relative to dendritic polarization (MedAPE: 6.54 – 15.73% M1–SO and 7.83 – 16.62% 4×1 HD axonal; 3.85 – 7.62% M1–SO and 4.42 – 8.42% 4×1 HD basal dendritic; 3.81 – 4.82% M1–SO and 4.6 – 5.3% 4×1 HD apical dendritic).

**Figure 7.**
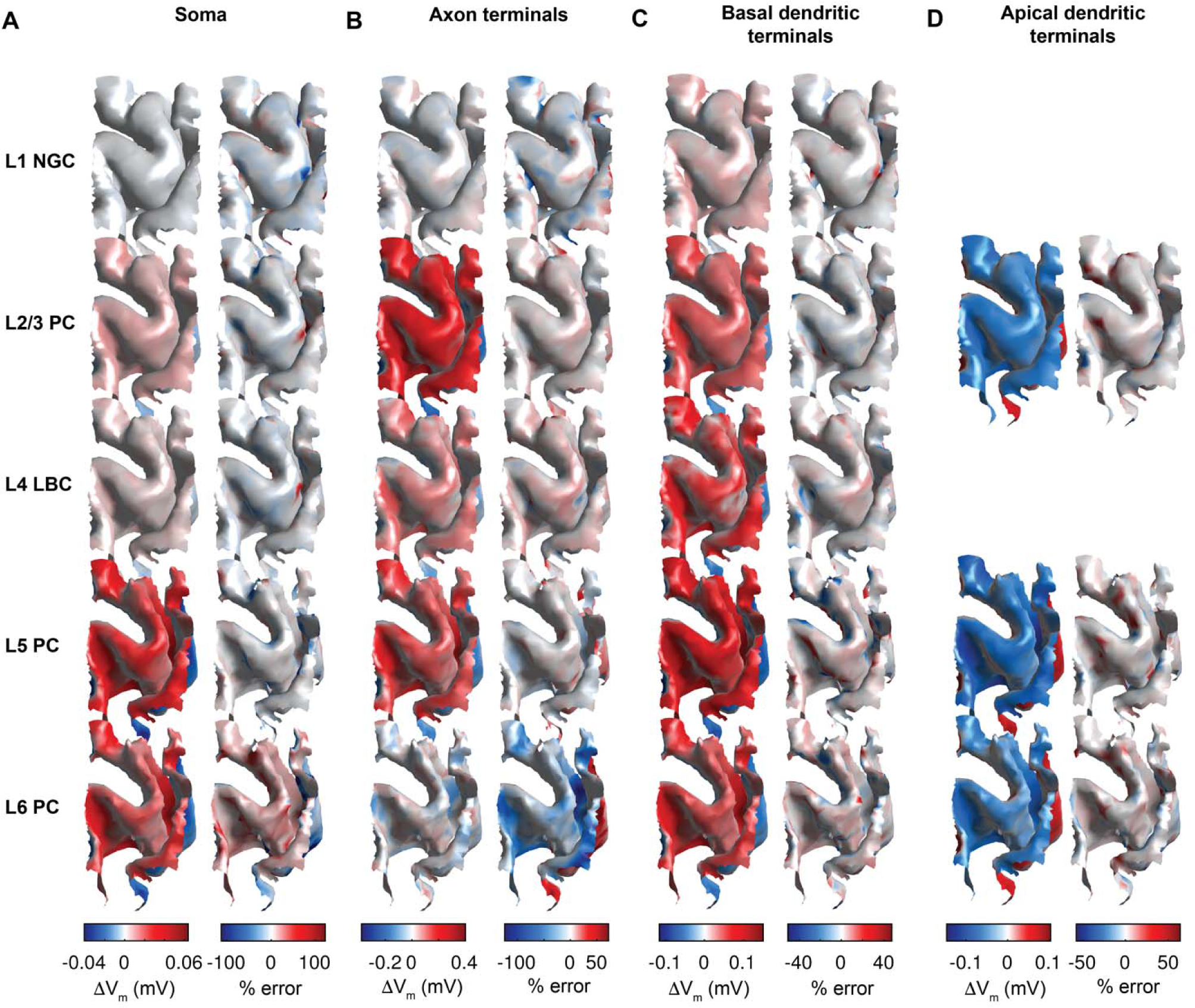
Spatial distribution of estimated polarization and error of tDCS generated polarizations using uniform E-field for M1–SO 7×5 cm rectangular pad montage at 1.8 mA current. Surface plots colored by polarization (left columns) and percent difference (right columns) of A) somatic polarization or peak polarization in B) axonal, C) basal dendritic, or D) apical dendritic terminal compartments when estimated using uniform E-field simulations as compared to FEM E-field coupled simulations at each location. Polarization value and percent difference at each point of surface plots is median across clones. Color bar limits set to 0.05 and 0.95 quantiles of percent error distributions across layers, which clips outliers due to extremely high percent error in regions of low polarization.

**Figure 8.**
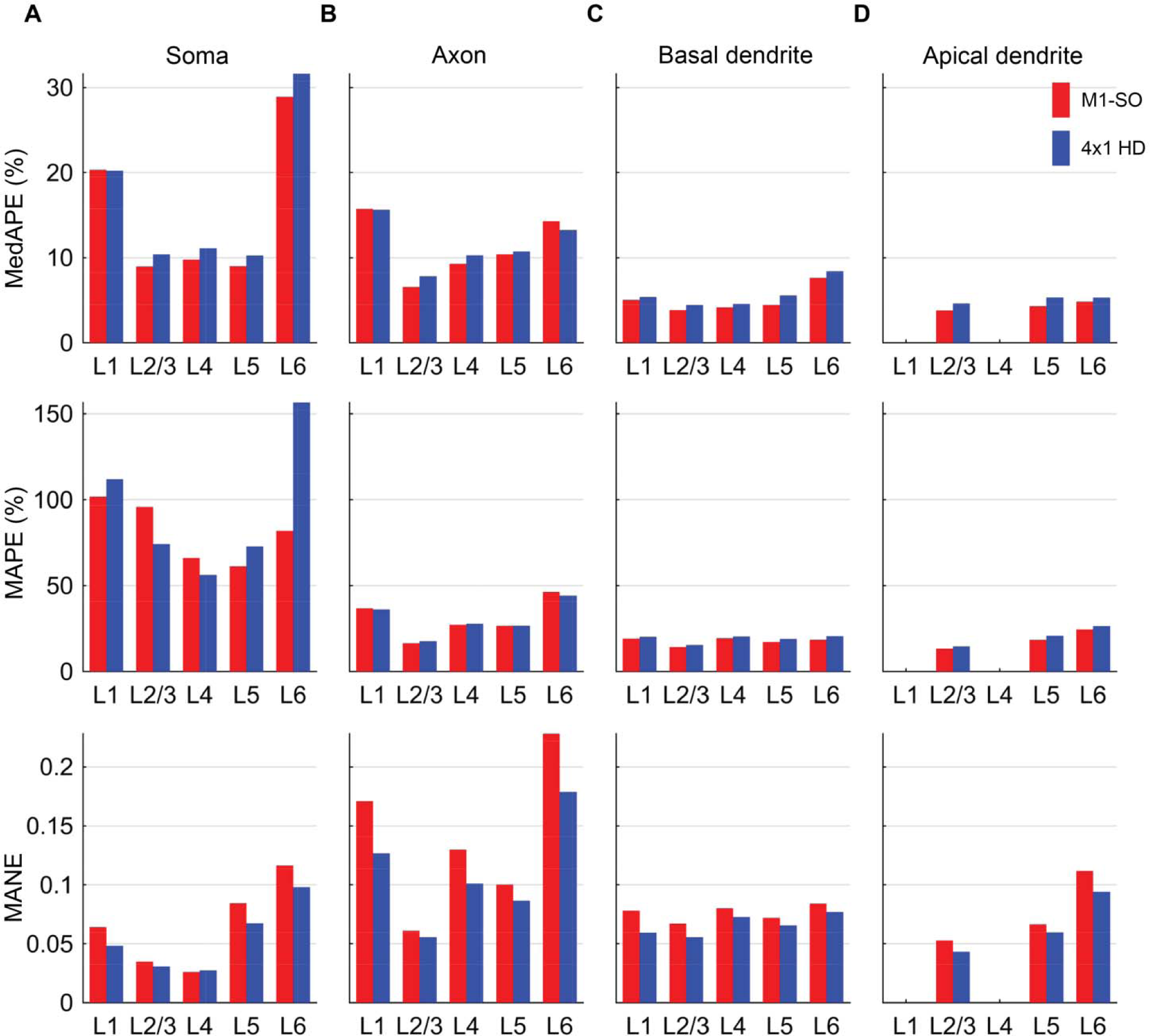
Layer-specific error statistics for estimated tDCS-generated polarization using uniform E-field for M1–SO 7×5 cm rectangular pad montage compared to polarizations simulated with FEM E-field. Bar plots indicate median absolute percent error (MedAPE) (top), mean absolute percent error (MAPE) (middle), and mean absolute normalized error (MANE) (bottom) for A) somatic polarization or peak polarization in B) axon terminal, C) basal dendritic terminal, or D) apical dendritic terminal compartments.

## 5. Discussion

We simulated neuronal polarization generated by both uniform E-field and E-field distributions generated by two commonly used tDCS montages in a multi-scale model consisting of an FEM E-field model with MRI-derived head geometry and biophysically realistic cable models of layer-specific excitatory and inhibitory neurons. The uniform E-field simulations demonstrated that the polarization of different neural compartments (soma, axon, and dendrites) across E-field directions varied by cell type. In the multi-scale model, we found not only that both the conventional (7×5 cm M1–SO) and more focal (4×1 HD) montages produce depolarization and hyperpolarization beneath the anodal contact, but also that axonal and dendritic polarization was substantially stronger than somatic polarization, with highest polarization at the terminations. For stimulus intensities generating peak |E| of ∼0.5 V/m in the target region (M1 hand knob), somatic polarization did not exceed 0.1 mV, while peak axonal polarization reached nearly 1 mV in the L2/3 PC axons terminals. Maximal dendritic polarization was observed in the inhibitory large basket cells (∼0.2 mV), closely followed by the L5 PCs (0.17 mV). Pyramidal cell somatic polarization was strongly correlated with the normal component of the E-field, while the correlation between with the normal component of the E-field and peak axonal and dendritic polarization was more modest. However, polarization of all compartments of interneurons was less well explained by the E-field normal component than in the pyramidal cells, indicating this component is a less effective proxy of interneuronal polarization. The uniform E-field approximation method captured broadly the spatial distribution of polarization in each compartment, but exhibited regions of high relative error, suggesting that capturing the non-uniform E-field distribution in the vicinity of individual neurons may be necessary for more accurate results, particularly in distal axonal compartments.

### 5.1. Implications for stimulation polarity and focality

A common heuristic in tDCS studies treats anodal tDCS as facilitatory and cathodal as inhibitory based on models and *in vitro* experiments showing inward current depolarizes and outward current hyperpolarizes the soma of pyramidal cells. These cellular effects are thought to explain the polarity-specific facilitation and inhibition of MEPs by anodal and cathodal tDCS, respectively, with the classical M1–SO montage [50]. However, our results illustrate that the distribution of polarization is more complex when considering detailed gyral anatomy in humans and morphological differences between cell types. Both the conventional and more focal electrode montages produced significant depolarization and hyperpolarization beneath the anode, with strongly bimodal distributions of peak polarization in the dendritic and axonal compartments. As expected from uniform E-field stimulations [18,51], apical dendritic polarization was always opposite in polarity to basal dendritic polarization in the pyramidal cells. Somatic polarization in the pyramidal cells largely followed the expected distribution, correlating strongly with the normal E-field component, although with the important caveat that cortical gyrification leads to both directions of current flow in the vicinity of the anode and cathode for even the more focal HD montage. Therefore, our results suggest that polarity-specific effects of tDCS on excitability may arise, instead, from more subtle shifts in balance of depolarization and hyperpolarization that occur with the change in stimulus polarity, or reversal of polarization polarity in specific populations of neurons, rather than a more uniform de- or hyperpolarization of the target region. In light of these results, it perhaps should be less surprising that the effects of stimulation polarity on neuromodulatory after-effects are more complex than the heuristic mentioned above implies. For example, Batsikadze et al. found that 20 min of 2 mA cathodal tDCS enhanced, rather than suppressed, MEP amplitude, while 1 mA cathodal stimulation produced the expected suppression [52]. In contrast, Ahn and Frohlich demonstrated consistent enhancement and suppression of MEP amplitude with 10 min of 2 mA anodal and cathodal tDCS, respectively [9].

The 4×1 HD montage was designed to produce more focal polarization, reducing current in off-target structures. One study comparing 4×1 ring HD and conventional 7×5 cm rectangular pad tDCS of M1 found stronger, longer-lasting effects with the HD montage in 14 subjects [53], but more replication is necessary to confirm the effect of focality on after-effects. Based on the simulated distributions of E-field, HD tDCS is clearly more focal on the global level, with peak E-field concentrated within the ROI encompassing the M1 hand region and the opposing postcentral gyrus, while the 7×5 cm rectangular pad electrodes generated peak E-field outside the region. With current intensities set to match peak E-field magnitude within the ROI, the distributions of polarization within each compartment were similar between the two tDCS montages. However, there were differences when analyzing polarization within sub-regions of the ROI, with the HD montage producing stronger polarization in the gyral crown and the M1–SO montage producing stronger polarization in the central sulcus. Still, the broad spatial distributions of polarization, especially in axonal and dendritic terminals, suggests tDCS may not produce specific effects based on the spatial focality of polarization alone; instead, specific effects may arise from selective modulation of active circuits or biasing of different synaptic inputs, as proposed by Bikson and Rahman [54].

### 5.2. Stimulation directionality

The local E-field direction determines the direction and magnitude of polarization; thus, a key question is how these direction-dependent effects relate to the subsequent neuromodulatory aftereffects. The importance of E-field direction relative to the neuronal elements in cortex has received recent interest as a way to extrapolate the neural effects from the electric field simulated in volume conductor models [13,55]. In particular, the normal component relative to the cortical surface is expected to correlate with somatic polarization of pyramidal cells and potentially to acute and lasting changes in excitability. Indeed, Laakso et al. found 20 min of 1 mA anodal M1–SO tDCS caused stronger suppression of MEP amplitudes in subjects with stronger *E_n_* in the M1 hand region, based on individualized head models. Rawji et al. applied bipolar tDCS to the M1 hand region in different orientations, finding posterior–anterior currents, but not anterior–posterior or latero–medial, caused significant suppression of MEP amplitudes relative to sham [55].

Our results suggest the direct neural effects may not be captured by simple decomposition of the E-field alone. We found the directional sensitivity of cell bodies, axons, and dendrites varies between excitatory pyramidal cells and inhibitory interneuronal cells. Pyramidal cell somatic polarization was indeed highly correlated with the E-field component normal to the cortical surface, but peak axonal and dendritic terminal polarization had more variable dependency on E-field direction due to their diffuse branching, which can also be seen directly in their polarization–direction maps (Supplementary Figure S2–S4). Furthermore, the apical dendrites of the pyramidal cells were always polarized in the opposite direction of the soma, i.e., inward currents depolarizing the soma concomitantly hyperpolarized the distal apical dendritic branches.

Peak axon terminal polarization was not well correlated with the tangential current, including of the interneurons, which are often considered to be “horizontally projecting” in contrast to radially directed pyramidal cells. Similarly, Rahman et al. found tangentially directed E-field had pathway-specific effects on synaptic efficacy depending on E-field orientation, but they found no average effect due to the range of pathway directions present within the slice [13]. Finally, while the interneurons experienced weak somatic polarization, the strongest dendritic polarization was generated in the L4 LBCs, raising the possibility tDCS could alter postsynaptic integration of inhibitory basket cells. Our results thus support the use of the normal E-field component as a proxy for pyramidal cell somatic polarization, but not for other cell types or neural elements. Experiments combining tDCS with subject-specific head models and brain recordings, rather than MEP measurements alone, will be necessary to further unravel the link between specific E-field components and neuromodulatory aftereffects.

### 5.3. Presynaptic mechanisms

One major finding of this study is the significantly stronger polarization of axon terminals relative to somatic polarization, reaching nearly an order of magnitude difference in the L2/3 PCs. The higher sensitivity of axon terminals to uniform E-field stimulation has been hypothesized based on cable theory [13,20,23,24] and demonstrated experimentally with patch-clamp recordings from L5 PC axon cut-endings (blebs) [21]. We quantified axon terminal polarization in models of both pyramidal and inhibitory cell types, accounting for their complex and cell-type specific local axon morphologies. Axon terminal polarization may recruit analog properties of action potential signaling in axons and presynaptic boutons revealed by axon bleb recordings and voltage imaging experiments [56–58]. Subthreshold presynaptic depolarization or hyperpolarization can modulate the amplitude of synaptic release, likely through changes in action potential waveform leading to altered calcium influx and neurotransmitter release [59]. Indeed, Chakraborty et al. found changes in action potential waveform properties in the axon bleb, but not soma, with 5 V/m uniform E-field stimulation [21]. In a computational model reproducing depolarization and hyperpolarization facilitation of synaptic transmission, Zbili et al. estimated ∼9 mV depolarization in a presynaptic bouton led to a 37% increase in synaptic transmission for a single AP [60]. Extrapolating from the peak axon polarizations in this study would suggest tDCS could produce up to ∼4% facilitation in L2/3 PCs and ∼1% facilitation in L4 LBCs axon terminals. While endogenous subthreshold polarization declines with distance from the cell body, tDCS produces maximal depolarization at distal axon terminals, with polarization proportional to branch length [24]. Experiments in cortical and hippocampal slices found evidence of acute changes in synaptic efficacy that are strongest with E-field aligned with the activated axonal inputs [13,25]. Using slices from motor cortex, Vasu and Kaphzan reported changes in the rate of spontaneous vesicle release by dc stimulation, dependent on voltage-gated sodium, potassium, and calcium channels [61–63], suggesting modulation of presynaptic release machinery. However, electrophysiological measures of synaptic transmission include the influence of postsynaptic mechanisms, and the effects of dc stimulation on action potential-evoked presynaptic release have yet to be measured. Our results suggest polarization of both excitatory and inhibitory axon terminals may play a role in the acute effects of tDCS. Future experimental work should characterize the effects of tDCS on presynaptic release of both excitatory and inhibitory cell types, as most work has focused on the post-synaptic compartments of pyramidal cells [13–15,18,22,64].

### 5.4. Comparison to previous work

Thus far, two other modeling studies coupled an FEM model of the E-field in the head to populations of spatially extended neuron morphologies to explore the effects of tDCS [33,65]. Seo et al. used models of a stellate cell, L3 PC, and L5 PC with idealized, straight axons that included a single terminal. They found strong correlation between pyramidal neuron somatic polarization and the E-field normal component, in agreement with our results. However, without realistic axon morphologies, they could not quantify the distribution of polarization within cell-type specific axon arbors as done in this study. Otherwise, several computational studies modeled uniform E-field stimulation with dc or ac waveforms in single neuron models with realistic morphologies [13,24,48], but these were limited to single pyramidal cell models. Another key advance is our inclusion of myelin in the axonal arbors, which increases the axonal length constant and coupling to the E-field, while reducing attenuation from terminals to more proximal axon compartments. More recently, Chung et al. simulated polarization and activation thresholds of one clone from the L2/3 PC, L4 LBC, and L5 PC models used in this study, based on models released with our previous publication [20], but did not analyze the relationship between subcellular polarization distributions and E-field components [33].

To study recruitment by tDCS of synaptic plasticity mechanisms, Kronberg et al. incorporated a phenomenological voltage-based model of long term potentiation and depression in a biophysically realistic hippocampal CA1 PC model that recapitulated measured effects of uniform dc E-field on LTP [15]. Subthreshold membrane polarization during a theta burst plasticity induction protocol enhanced or suppressed the magnitude of LTP depending on stimulus polarity, and these effects were correlated with changes in firing probability due to somatic or dendritic polarization. More recent results from the same group suggest tDCS-induced changes in LTP may be mediated by increased somatic firing from both direct somatic polarization and increased network activity [64]. However, this model did not include distal axonal compartments of the modeled cell or presynaptic inputs, precluding analysis of axon-dependent mechanisms. Additionally, it is worth noting these experiments were conducted with 20 V/m E-field amplitudes, around 20–40× higher than typical tDCS peak E-fields. While it is possible that these same phenomena are recruited with lower E-field magnitudes, the relative recruitment of pre- and postsynaptic mechanisms at realistic field amplitudes may be significantly different. Ultimately, developing models combining dendritic and axonal morphologies with mechanisms of both post-synaptic and pre-synaptic plasticity will be crucial for identifying the relative contributions of polarization at different subcellular compartments to the neuromodulatory effects of tDCS.

### 5.5. Quasi-uniform assumption

Using uniform E-field to approximate the non-uniform tDCS generated E-field distributions for each neuron in the ROI produced distributions of polarization that matched qualitatively the distributions with the full, non-uniform E-field, while also yielding regions with substantial error. The estimated peak axonal and dendritic polarization even had the incorrect polarity at some positions. These errors were predominantly a result of the E-field at the most polarized element (e.g., axon or dendritic terminal) differing in magnitude than the E-field at the soma. In contrast, the E-field direction did not vary significantly within single morphologies. Rather than producing gradients high enough to polarize processes directly via the second difference (“activating function”) [66], the E-field non-uniformity leads to varying E-field magnitude at the different morphological discontinuities, where polarization is proportional to the E-field magnitude [20,24]. Thus, the imposed transmembrane currents along the morphology in the non-uniform E-field differ from a uniform E-field with matched intensity and direction at a single location. Finally, axial current redistribution within the morphology leads to further differences in the final steady-state polarization in each compartment than would be expected given just the local extracellular E-field [67]. Still, errors for peak axonal and dendritic polarization estimation would likely be reduced if the E-field were extracted at the specific compartments of interest, allowing a more local (compartment-specific) approximation of the E-field as uniform. Using uniform E-field in *in vitro* studies will remain a useful way of approximating transcranial stimulation modalities while controlling for E-field direction, particularly for studying effects at a single location within the neuron (e.g., soma or dendrites). However, our results suggest accurate modeling of distributions of transmembrane polarization across the entire spatial extent of cortical neurons, especially in the axon arbor, requires modeling the tDCS current flow in realistic geometries.

We also found that estimating polarization based on the uniform E-field at the soma resulted in large errors of the predicted polarization, especially in the axons. Therefore, further work developing accurate methods of efficiently approximating the response of biophysically realistic cortical neurons will be important to making these models more broadly accessible, and it will have to consider the effect of non-uniform E-fields across the neuron morphology. We recently showed 3D convolutional neural networks can accurately predict activation thresholds based on sparse samples of the local E-field distribution [68], suggesting similar approaches may allow rapid estimation of neuronal polarization by tDCS.

### 5.6. Limitations

As noted previously [32], this model does not incorporate medium or long range afferent fibers, which we expect would also experience significant polarization at their terminals depending on their orientation within the E-field. Incorporating these axons as they traverse gray and white matter with microscopic detail is a significant challenge; recently developed methods for imaging high-resolution volumetric tissue samples, segmenting neural morphologies [69–71], and reconstructing full axons of single neurons [72] may provide data sources to incorporate these elements in multi-scale models. We also focused here on a single azimuthal rotation of each model neuron, but our new method for fitting neuron morphologies within the cortical geometry also generates other valid azimuthal rotations, which are included in the shared code.

Furthermore, the scope of this study was limited to quantifying direct polarization of cortical neurons, but a key next step will be to test how these direct effects alter endogenous activity at the single cell and network levels. The effect of tDCS on single neuron firing and action potential propagation can be readily explored in these models by simulating different firing patterns with synaptic inputs distributed along the somatodendritic arbor or somatic current injection. Simulating large-scale networks with the full multi-compartmental models used in this study, especially including the axon, is computationally demanding. The full Blue Brain network model, from which our models were derived originally, required powerful supercomputers to simulate the activity of 31,000 neurons without their axons included [40]. For simulations on more accessible hardware, the cell-type and compartment-specific distributions of polarization computed in this study can be used in large scale network models of tDCS with simplified representations of individual neurons to study how effects on single neurons might alter network activity. Our results suggest network models with single compartment (point) neuron models may be neglecting relevant effects on presynaptic and postsynaptic compartments, although they may lend insight into the mechanisms by which somatic polarization could alter network activity [73].

Finally, while our models captured the realistic morphology of local axon collaterals, the specific properties and composition of ion channels in distal axons are still unclear due to their inaccessibility to electrical recording techniques. We modeled the axon terminals with similar properties to the nodes of Ranvier, which is likely an oversimplification. Recent patch clamp recordings in cultured neocortical presynaptic boutons suggest they have high densities of sodium, potassium, and calcium channels, leading to large, stable action potentials [74]. Additionally, presynaptic boutons and synaptic clefts have unique ultrastructure, with specialized cytoarchitecture for trafficking and exocytosis of synaptic vesicles at the active zone on the intracellular side [75,76] and tightly packed pre- and postsynaptic membrane (15–25 nm apart [77]) as well as astrocytic processes [78] on the extracellular side. The geometric and electrical properties of these structures may influence current flow or extracellular voltage at synapses and hence impact terminal polarization. Models that capture the electrophysiological properties of synaptic boutons, including mechanisms of neurotransmitter release, will be necessary to characterize fully the acute and lasting effects of tDCS on presynaptic terminals. Besides axon terminals, tDCS may also affect non-neuronal cells, such as glia [79] and vasculature [80], which were not captured in our model.

## 6. Conclusion

Coupled E-field–neuron simulations of tDCS enable calculation of the direct neural polarization and capture the specific electrode placement, geometry, and temporal waveform. Our results support the importance of extending E-field models to include acute representations of neurons, as exemplified by the diverse responses of different cortical cell types and subcellular elements. While the uniform E-field approximation can lend significant insight, the non-uniform E-field distribution generated along single neurons by tDCS produces different distributions of steady-state polarization than predicted by uniform E-field stimulation. Finally, tDCS generated the strongest polarization at the axonal and dendritic terminals, with substantially weaker somatic polarization in all cell types, providing a putative mechanism by which both excitatory and inhibitory cell types may be modulated directly by tDCS with subthreshold intensities.

## 7. Acknowledgements

Preliminary results from this work were presented at the NYC Neuromodulation Conference and NANS Summer Series in 2017, the NYC Neuromodulation 2020 Online Conference, and the 2021 Society for Neuroscience Conference. We thank Adrian Lopez, Dr. Moritz Dannhauer, and Dr. Boshuo Wang for their technical assistance and helpful discussions. We also thank the Duke Compute Cluster team for computational support. Research reported in this publication was supported by the National Institutes of Health under Award Numbers R01NS088674, R01NS117405, and R01MH128422 as well as by the National Science Foundation under Graduate Research Fellowship DGF 1106401. The content is solely the responsibility of the authors and does not necessarily represent the official views of the funding agencies.

## 8. Declaration of Interest

A.V.P. is an inventor on patents on TMS technology, has equity options and serves on the scientific advisory board of Ampa, and has received research funding, travel support, patent royalties, consulting fees, equipment loans, hardware donations, and/or patent application support from Rogue Research, Magstim, MagVenture, Neuronetics, BTL Industries, Magnetic Tides, Soterix Medical, and Ampa. The other authors declare that they have no known competing financial interests or personal relationships that could have appeared to influence the work reported in this paper.

## Notes

### Competing Interest Statement

The authors have declared no competing interest.

