## Supplementary Materials for "Multi-scale model of axonal and dendritic polarization by transcranial direct current stimulation in realistic head geometry"

**Supplementary Material**

Authors: Aman S. Aberra*^1^, Ruochen Wang^1,2^, Warren M. Grill^1,3,4,5^ Angel V. Peterchev^1,2,3,5^

^1^Dept. of Biomedical Engineering, School of Engineering, Duke University, NC

^2^Dept. of Psychiatry and Behavioral Sciences, School of Medicine, Duke University, NC

^3^Dept. of Electrical and Computer Engineering, School of Engineering, Duke University, NC

^4^Dept. of Neurobiology, School of Medicine, Duke University, NC

^5^Dept. of Neurosurgery, School of Medicine, Duke University, NC

*Present affiliation: Dept. of Biological Sciences, Dartmouth College, Hanover, NH

Email addresses:

, and

Corresponding author: A.V. Peterchev

Address: Department of Psychiatry and Behavioral Sciences, Duke University Medical Center,

Box 3620, DUMC, Durham, NC 27710, USA

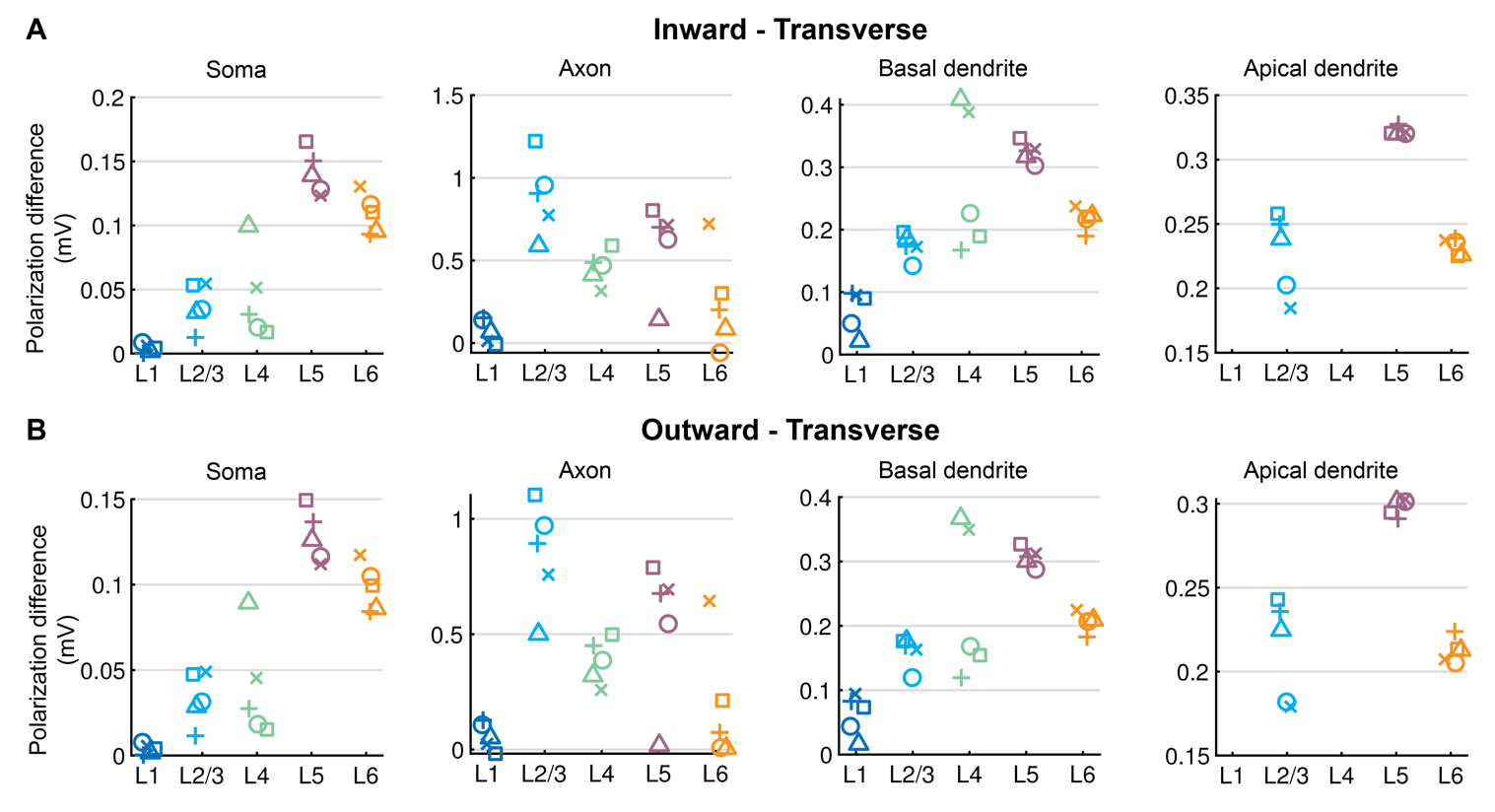

Supplementary Figure S1. Polarization differences for transverse and inward/outward uniform E-field. Difference in mean peak polarization for A) inward versus transverse E-field and B) outward versus transverse E-field in somatic, axonal, basal dendritic, and apical dendritic compartments. Mean peak polarizations computed for E-fields directed transverse ($\boldsymbol{60^{\circ}<\theta\leq120^{\circ}}$) and approximately parallel to the somato–dendritic axis—either upwards, towards the pial surface ($\boldsymbol{0\leq\theta\leq60^{\circ}}$), or downwards, towards white matter ($\boldsymbol{120^{\circ}<\theta\leq180^{\circ}}$)—after averaging peak polarizations across all azimuthal rotations within the relevant polar angles. Symbols for each clone match Figure 2E.

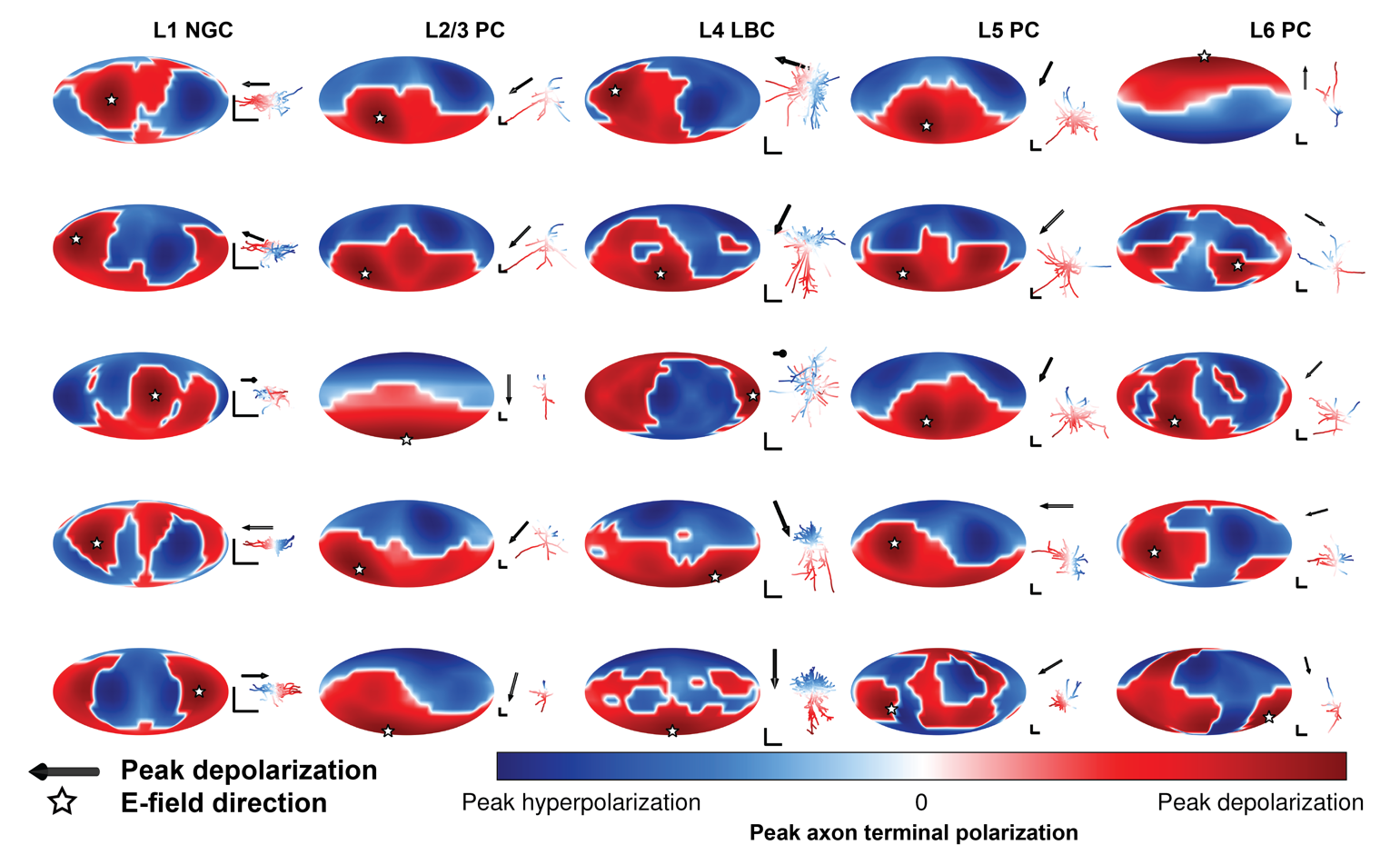

Supplementary Figure S2. Polarization–direction maps of peak axon terminal polarization for all cell types and their virtual clones. Peak axonal polarization always occurred at a terminal. As in Figure 2, clones of each cell type (columns) are ordered by peak axonal depolarization, with lowest peak depolarization at the bottom. See Figure 2E for absolute values of peak axonal polarization in mV for each clone. Corresponding axon morphology (excluding dendritic arbor) plotted to the right of each map colored with polarization distribution for E-field direction producing peak axon terminal depolarization, indicated by black arrow and matching white star. All scale bars are 250 µm.

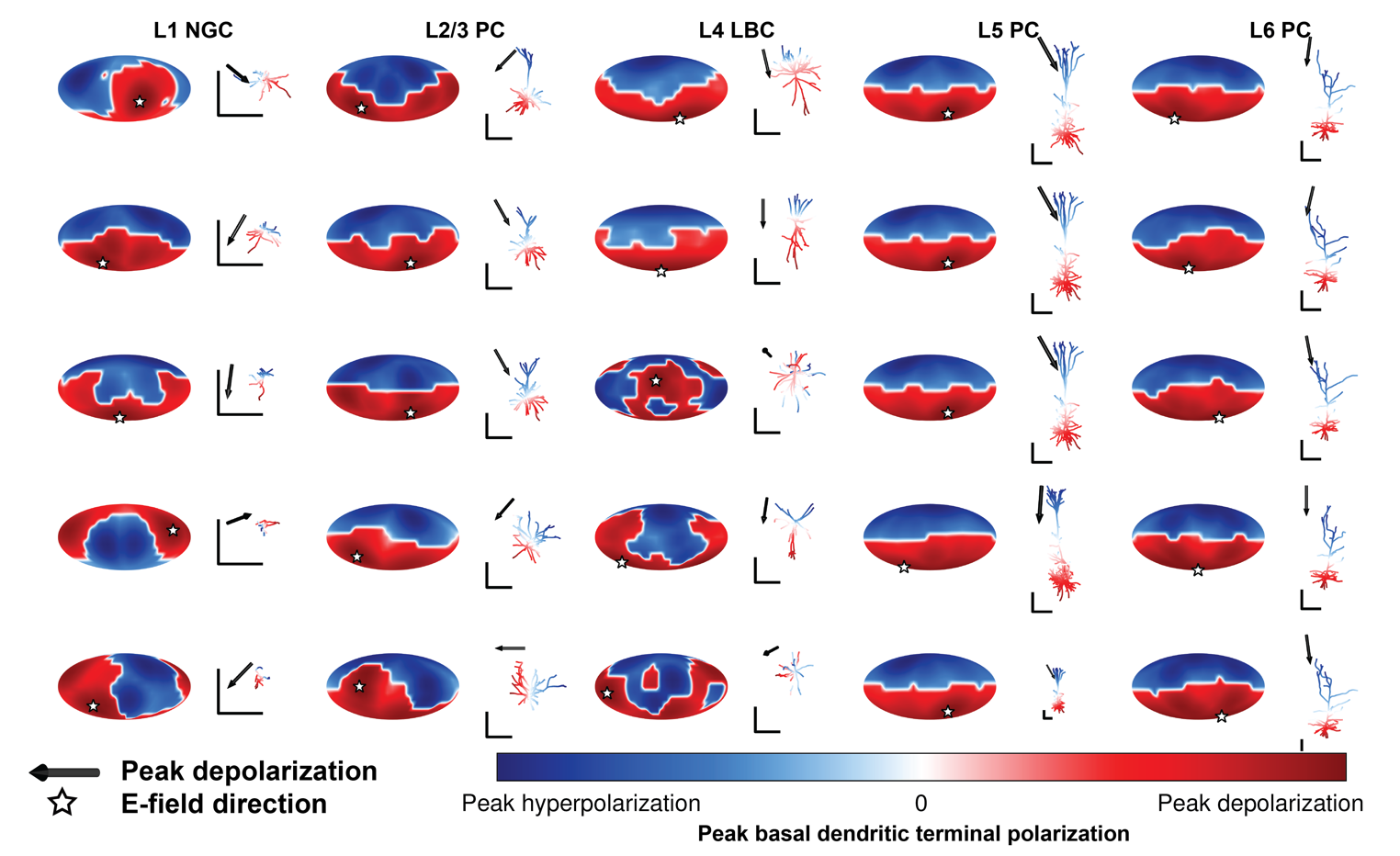

Supplementary Figure S3. Polarization–direction maps of peak basal dendritic terminal polarization for all cell types and their virtual clones. Peak basal dendritic polarization always occurred at a terminal. As in Figure 2, clones of each cell type (columns) are ordered by peak basal dendritic terminal depolarization, with lowest peak depolarization at the bottom. See Figure 2E for absolute values of peak axonal polarization in mV for each clone. Corresponding dendritic morphology (excluding axonal arbor) plotted to the right of each map colored with polarization distribution for E-field direction producing peak basal dendritic terminal depolarization, indicated by black arrow and matching white star. All scale bars are 250 µm.

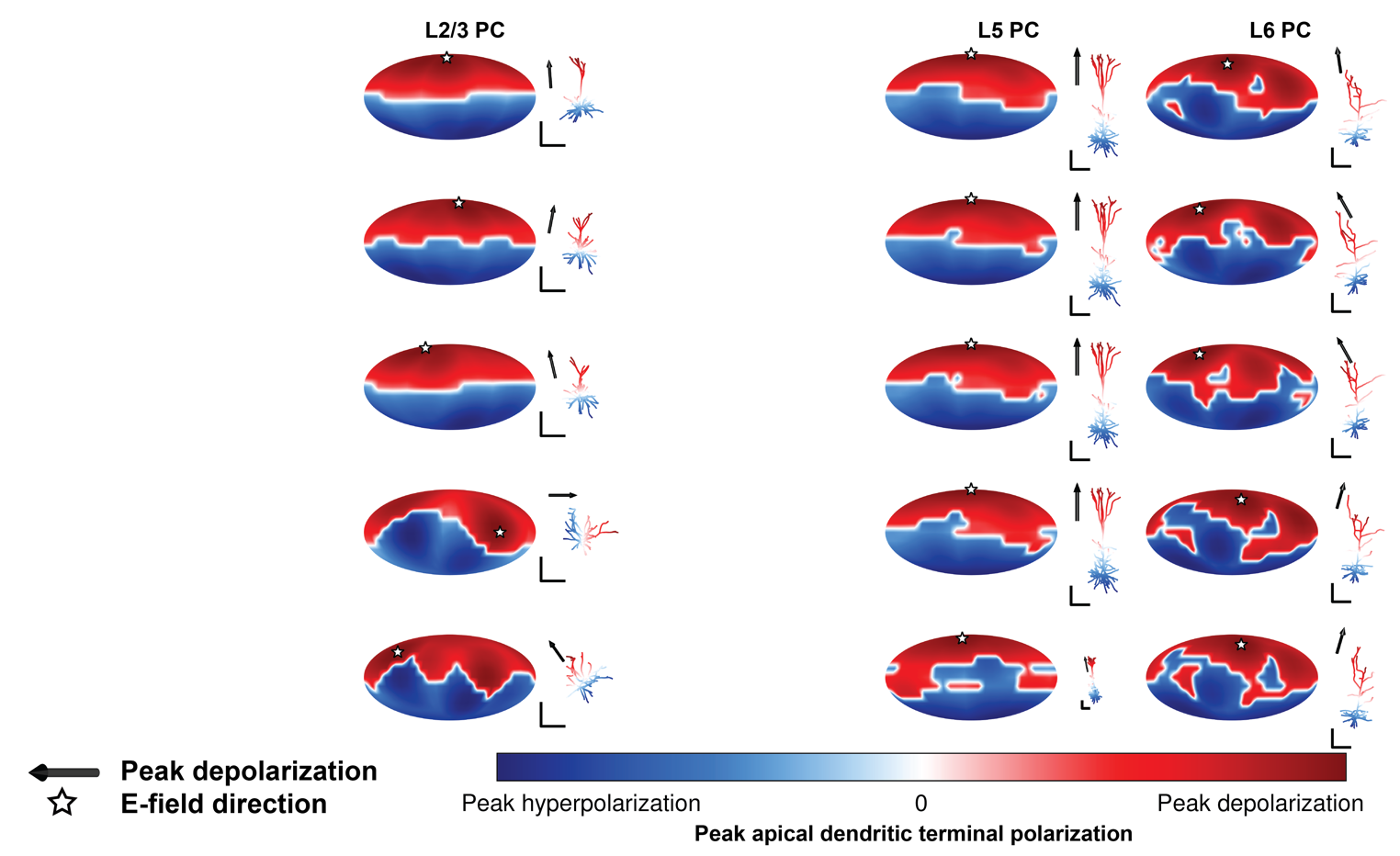

Supplementary Figure S4. Polarization–direction maps of peak apical dendritic terminal polarization for all cell types and their virtual clones. Peak apical dendritic polarization always occurred at a terminal. As in Figure 2, clones of each cell type (columns) are ordered by peak apical dendritic depolarization, with lowest peak depolarization at the bottom. See Figure 2E for absolute values of peak apical dendritic terminal polarization in mV for each clone. Corresponding dendritic morphology (excluding axonal arbor) plotted to the right of each map colored with polarization distribution for E-field direction producing peak apical dendritic terminal depolarization, indicated by black arrow and matching white star. All scale bars are 250 µm. L1 NGC and L4 LBC excluded due to lack of apical dendrites in these cell types.

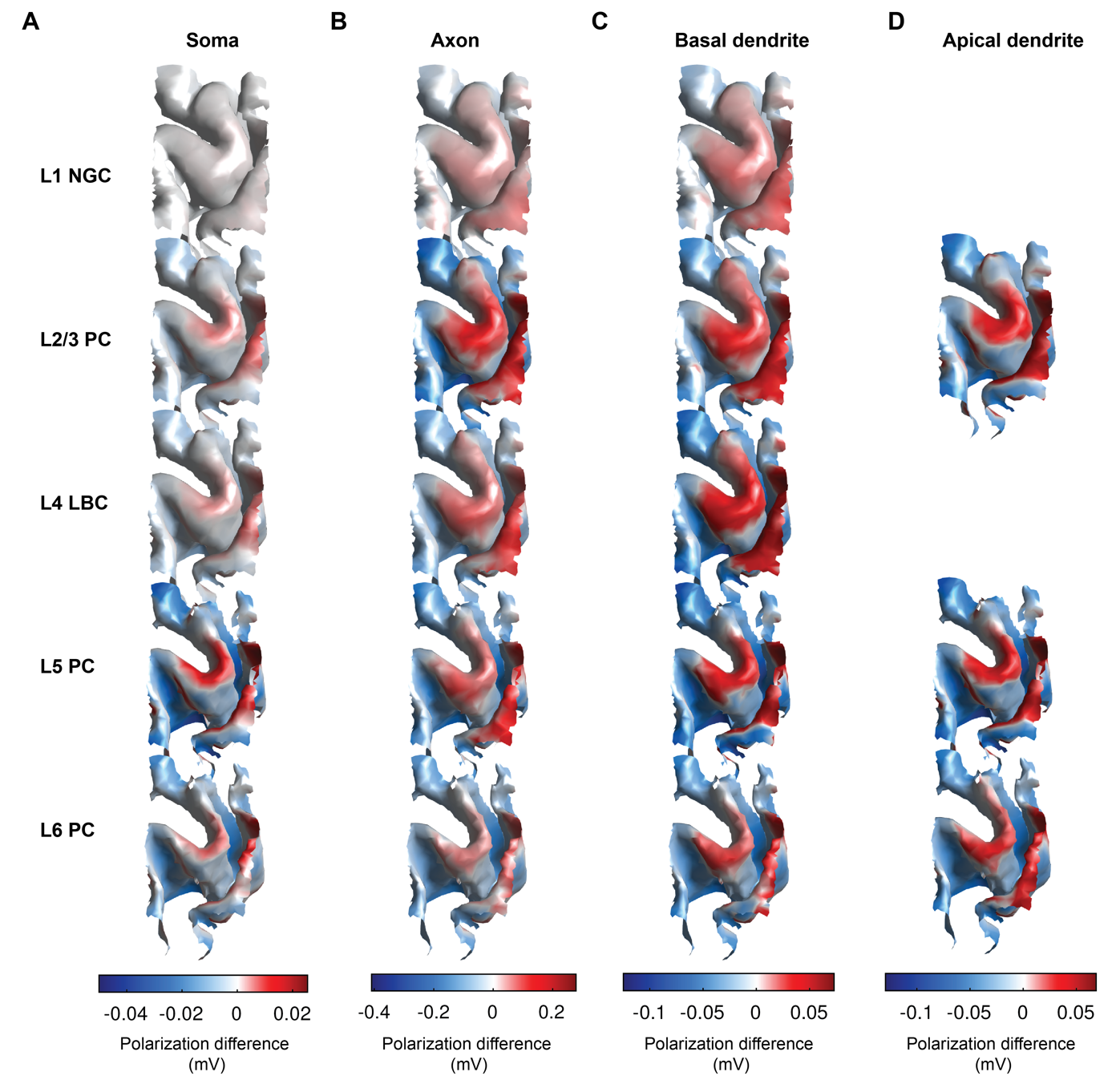

Supplementary Figure S5. Difference maps of absolute membrane polarization between M1–SO 7×5 cm rectangular pad and 4×1 HD tDCS in hand knob ROI (4×1 HD – M1–SO). Surface plots are colored by median difference across clones at each position of A) somatic polarization, or peak polarization in B) axonal, C) basal dendritic, and D) apical dendritic compartments. Positive values (red) indicate stronger polarization generated by 4×1 HD and negative values (blue) indicate stronger polarization generated by M1–SO montage.

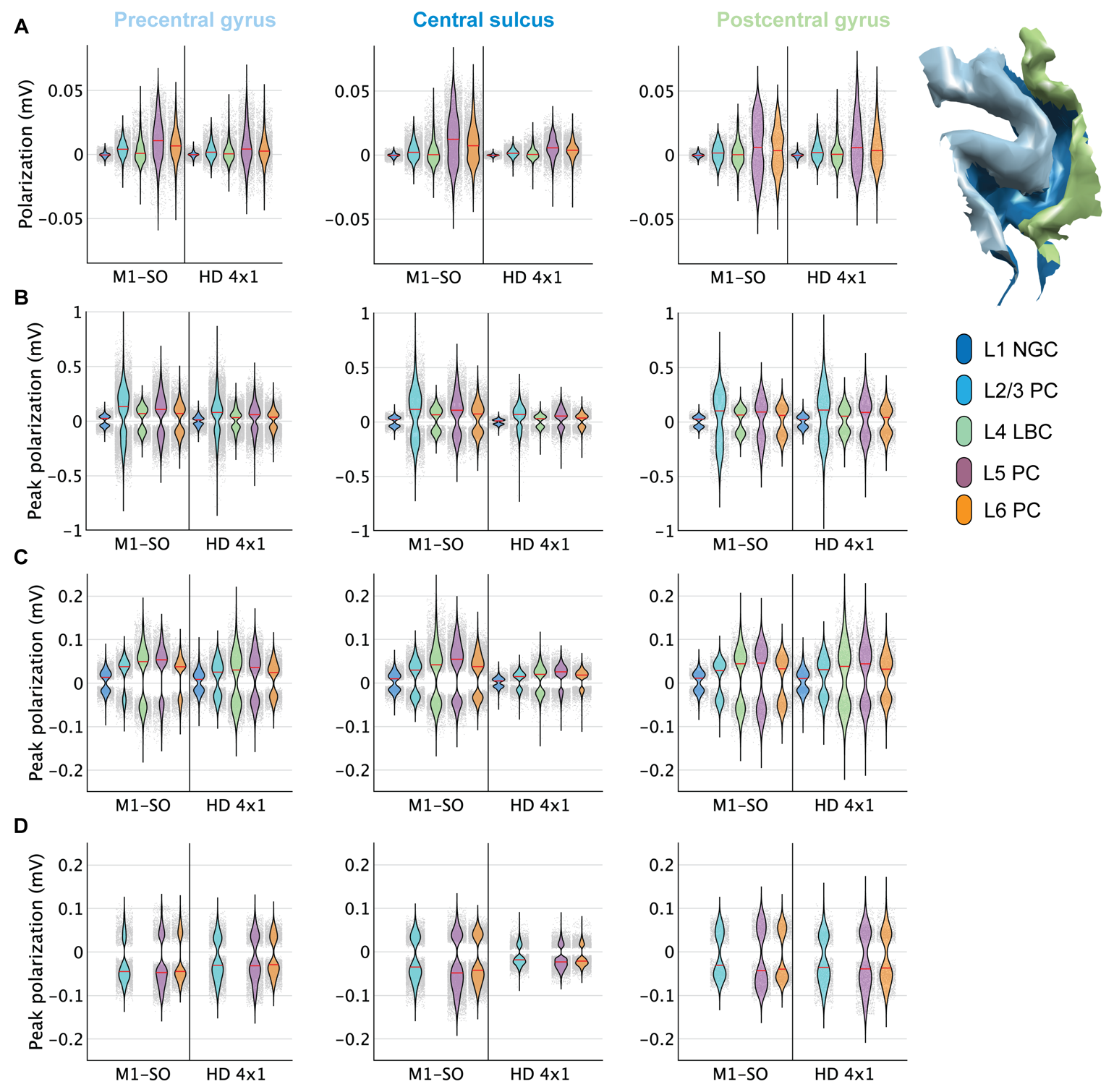

Supplementary Figure S6. Polarization distributions in ROI with M1–SO 7×5 cm rectangular pad and 4×1 HD tDCS, divided by atlas-defined cortical region. Each violin plot depicts the probability density estimate of the A) somatic polarization, or peak polarization in B) axonal, C) basal dendritic, and D) apical dendritic compartments of each layer for both electrode montages, within the three sub-regions of the ROI: Precentral gyrus (left), central sulcus (middle), and postcentral gyrus (right). Probability density was estimated using a normal kernel smoothing function with optimal bandwidth for each distribution. Point spread of polarization values plotted in gray, with each point representing the polarization value from a single model neuron. Right: Three sub-regions shown on L5 surface mesh, labels derived from Freesurfer automatic cortical parcellation based on the Destrieux standardized atlas [42].

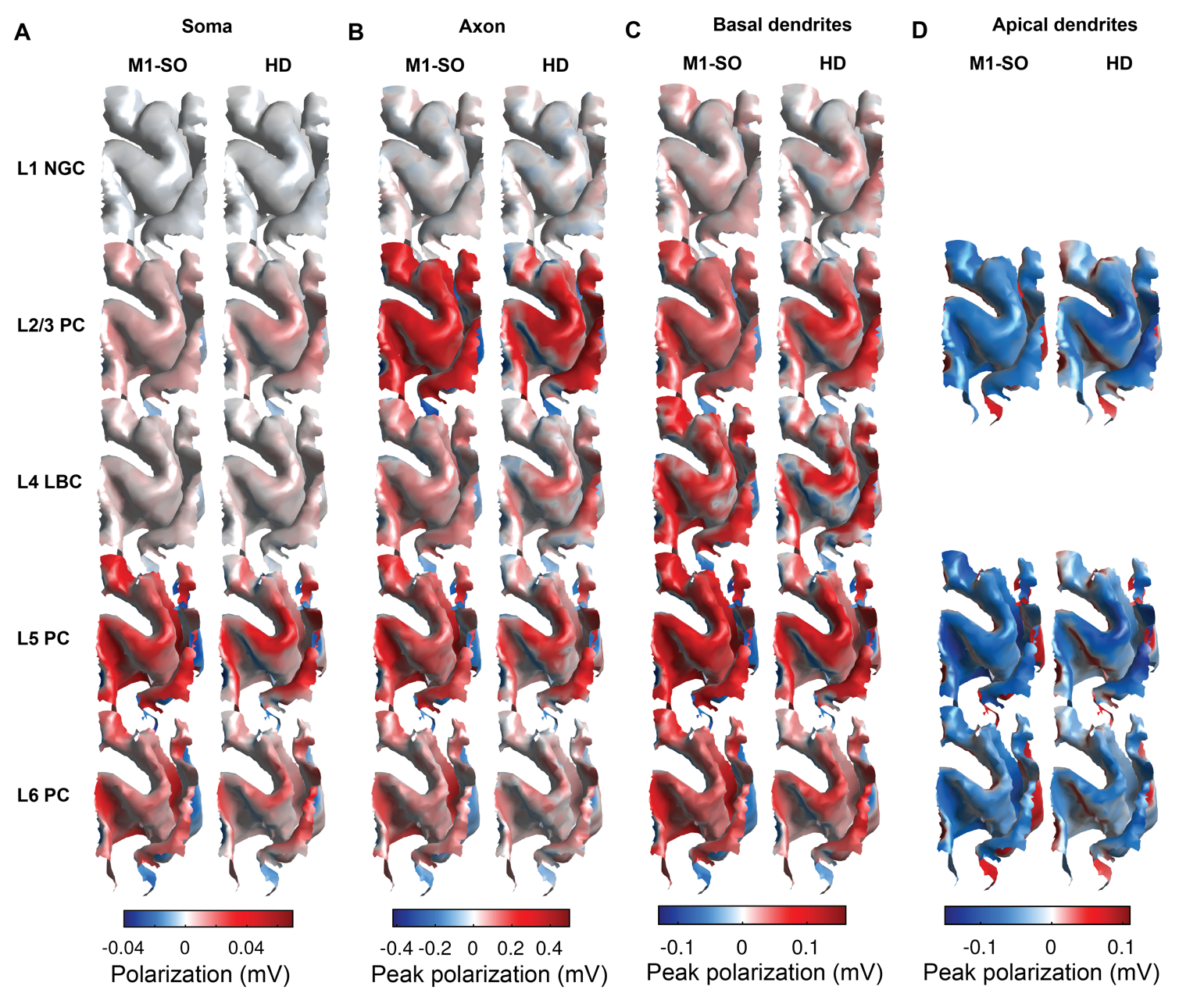

Supplementary Figure S7. Spatial distribution of membrane polarization in ROI using uniform E-field approximation method with M1–SO 7×5 cm rectangular pad and 4×1 HD tDCS. Surface plots are colored by median across clones at each position of A) somatic polarization, or peak polarization in B) axonal, C) basal dendritic, and D) apical dendritic compartments for 1.8 mA M1–SO rectangular pad (left) and 2.0 mA 4×1 HD tDCS.

|  | Soma | | | Peak axonal | | | Peak basal dendritic | | | Peak apical dendritic | | |
| --- | --- | --- | --- | --- | --- | --- | --- | --- | --- | --- | --- | --- |
| Montage | MedAPE (%) | MAPE (%) | MANE | MedAPE (%) | MAPE (%) | MANE | MedAPE (%) | MAPE (%) | MANE | MedAPE (%) | MAPE (%) | MANE |
| M1–SO | 12.93 | 81.31 | 0.12 | 10.66 | 30.65 | 0.16 | 4.80 | 17.71 | 0.09 | 4.29 | 18.74 | 0.11 |
| 4×1 HD | 14.10 | 94.36 | 0.09 | 11.17 | 30.45 | 0.13 | 5.46 | 19.03 | 0.07 | 5.04 | 20.57 | 0.10 |

Supplementary Table 1. Summary error statistics for estimated tDCS-generated polarization using uniform E-field across layers. MedAPE: Median absolute percent error; MAPE: Mean absolute percent error; MANE: Mean absolute normalized error.

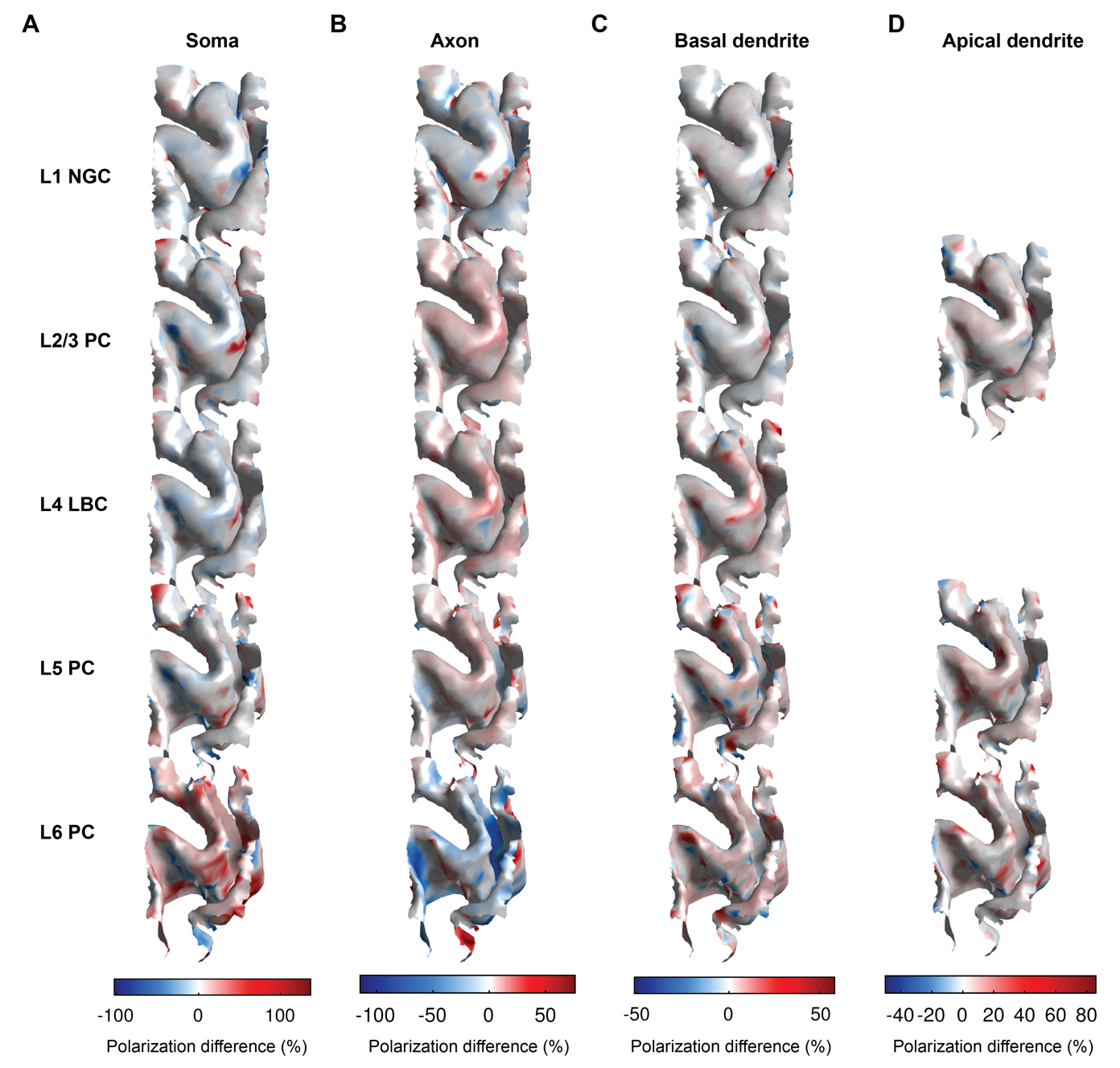

Supplementary Figure S8. Estimation error of tDCS generated polarizations using uniform E-field for 4×1 HD montage. Surface plots colored by percent difference of A) somatic polarization or peak polarization in B) axonal, C) basal dendritic, or D) apical dendritic compartments when estimated using uniform E-field simulations vs. FEM E-field coupled simulations at each location. Percent difference at each point of surface plots is median across clones. Color bar limits set to 0.05 and 0.95 quantiles of percent error distributions across layers, which clips outliers due to extremely large percent error in regions of low polarization.

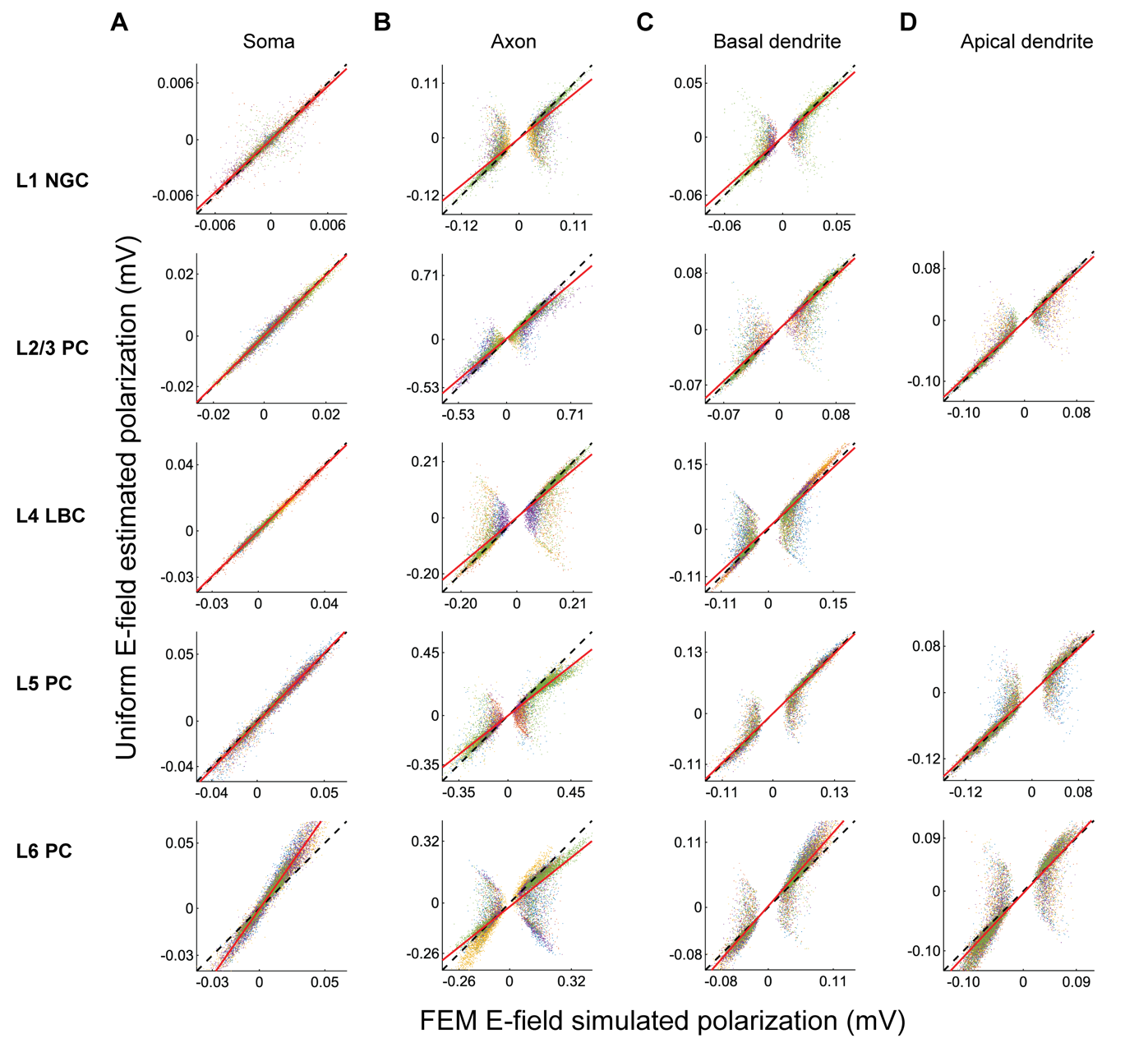

Supplementary Figure S9. Correlation of estimated tDCS generated polarization using uniform E-field for M1–SO 7×5 cm rectangular pad montage with polarizations simulated with FEM E-field. Each scatter plot includes simulated vs. estimated polarization value for each model neuron in the ROI, colored separately for each clone for A) somatic polarization or peak polarization in B) axonal, C) basal dendritic, or D) apical dendritic compartments. Also included are unity line (dashed black) and linear regression (solid red).
